# Robo2 drives target selective peripheral nerve regeneration in response to glia derived signals

**DOI:** 10.1101/2020.12.07.406942

**Authors:** Patricia L. Murphy, Jesse Isaacman-Beck, Michael Granato

**Affiliations:** Department of Cell and Developmental Biology, Perelman School of Medicine, University of Pennsylvania, Philadelphia, Pennsylvania, United States of America

**Keywords:** peripheral nerve regeneration, nerve branch, zebrafish, around-about receptor, *robo2*, Collagen4a5, target selection

## Abstract

Peripheral nerves are divided into multiple branches leading to divergent synaptic targets. This poses a remarkable challenge for regenerating axons as they select their original trajectory at nerve branch-points. Despite implications for functional regeneration, the molecular mechanisms underlying target selectivity are not well characterized. Zebrafish motor nerves are composed of a ventral and a dorsal branch that diverge at a choice point, and we have previously shown that regenerating axons faithfully select their original branch and targets. Here we identify Robo2 as a key regulator of target selective regeneration. We demonstrate that Robo2 function in regenerating axons is required and sufficient to drive target selective regeneration, and that Robo2 acts in response to glia located precisely where regenerating axons select the branch-specific trajectory to prevent and correct axonal errors. Combined our results reveal a glia derived mechanism that acts locally via axonal Robo2 to promote target selective regeneration.

## Introduction

The peripheral nervous system (PNS) has retained a remarkable capacity for axonal regeneration. Following injury, a well characterized injury response system transmits injury signals to the cell body initiating coordinated changes in cellular morphology, gene expression and chromatin remodeling (reviewed in ref. ^1^). Over the past decades molecular pathways that dramatically increase the rate of regenerative axonal regrowth have been identified ^2–5^. Moreover, supplementing pro-regenerative factors to regenerating PNS axons accelerates their regenerative growth rates, yet falls short of providing spatial information to direct axons towards their original, preinjury targets ^6–9^. While components of many developmental axon guidance pathways are upregulated after nerve injury, their functional roles in directing regenerating axons are largely unknown (reviewed in ref. ^10^) Thus, how regenerating axons navigate an environment that differs drastically from an embryonic environment, including how they select their original trajectories when confronted with nerve branch points is not well understood.

In vertebrates, individual peripheral nerves exiting from the spinal cord divide repeatedly into a series of progressively smaller branches, each carrying axons that innervate distinct synaptic targets ^11^. Depending on the type and location of a nerve injury, regenerating axons encounter a series of branch-points as they extend toward their original targets. Thus, for regenerating axons repeatedly selecting their appropriate branch at each branch-point, while a critical step to ensure functional regeneration, represents a formidable challenge. There is considerable evidence that axons have retained some capacity to select their original nerve branch ^12,13^, supporting the existence of dedicated molecular mechanisms to govern this process. Although branch-specific regeneration of peripheral axons was first demonstrated over 50 years ago ^14,15^, the molecular mechanisms that underly branch-selective, and hence target-selective regeneration are largely unknown. Given the complexity of the process it is maybe not surprising that the regenerative capacity to select the original nerve branch is limited, causing regenerating axons to frequently select inappropriate paths at nerve branch-points ^16,17^. For example, motor axons that inappropriately regenerate into nerve branches innervating either the skin or muscles antagonistic to their original target can significantly reduce the level of functional regeneration ^18,19^.

We have previously established the optically transparent larval zebrafish as model system to study the cellular and molecular mechanisms of target selective peripheral nerve regeneration *in vivo* ^20^. Zebrafish spinal motor nerves are composed of functionally distinct axonal populations that share a common path before diverging at a defined point into two major branches: a ventral nerve branch consisting of ~50 individual motor axons that innervate ventral muscle territories, and a dorsal nerve branch consisting of ~20 motor axons that innervate dorsal muscle territories ^21–24^. Following nerve transection regenerating spinal motor axons reliably choose the nerve branch that leads to their original muscle targets ^12,20^, and live cell imaging revealed that regenerating axons of the dorsal nerve branch pause at the nerve branch-point and pause, explore both the incorrect ventral and the original dorsal path before selecting their correct dorsal path. During this exploratory period, a small group of Schwann cells at the nerve branch-point simultaneously upregulates the expression of the extracellular matrix (ECM) component *col4a5* and the repulsive axon guidance cue *slit1a* ^12^, which have been shown to bind each other with high affinity ^25^. Moreover, we previously demonstrated that *col4a5* is required during regeneration to guide regenerating dorsal axons ^12^, and proposed a model by which Schwann cell-derived Col4a5 scaffolds Slit at the nerve branch-point to prevent regenerating dorsal nerve branch axons from inappropriately entering into and extending along inappropriate trajectories (Fig. 1A-D, ref ^12^).

**Fig. 1.**
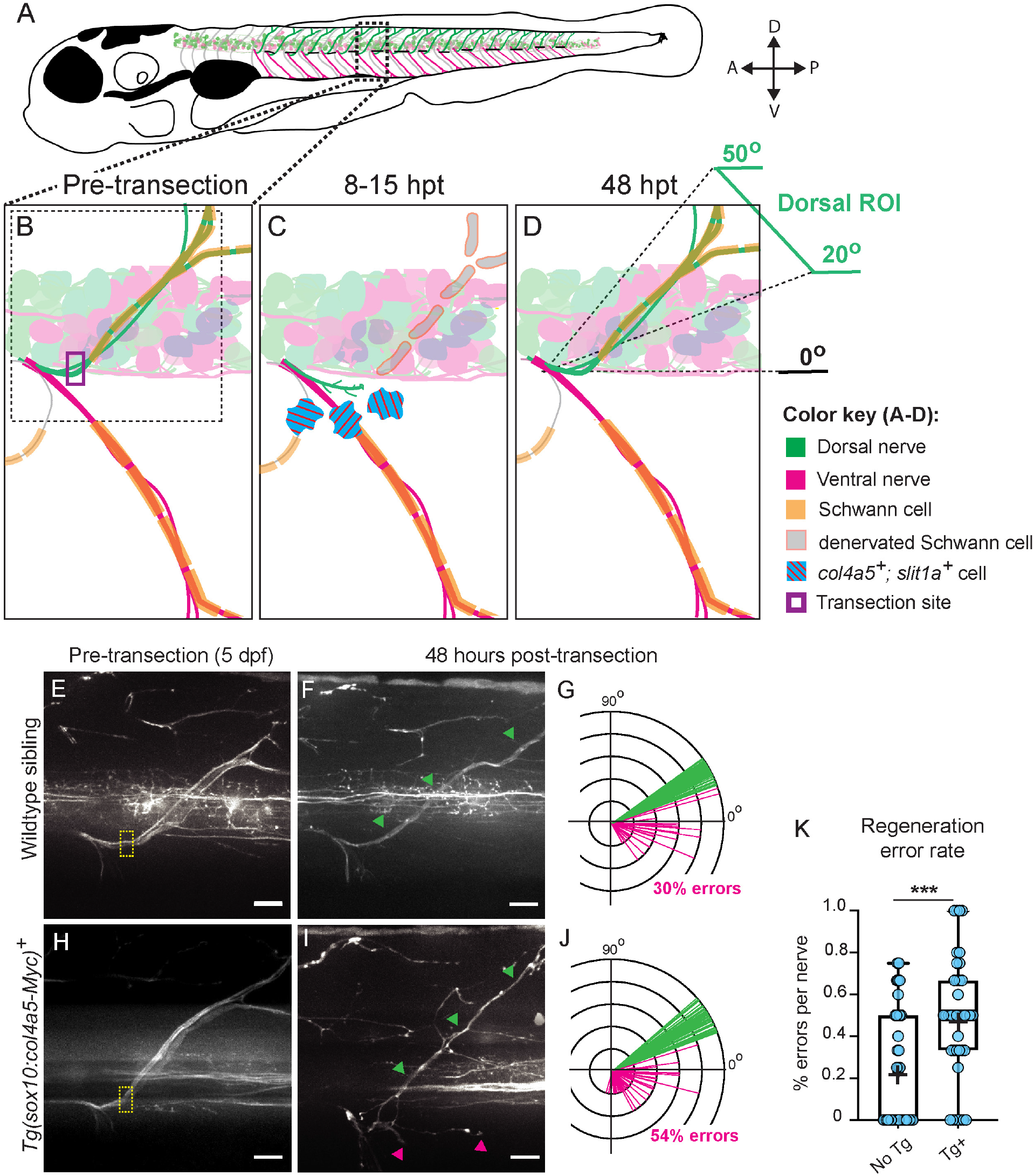
Spatiotemporal restriction of col4a5 to the nerve branch-point is required for target-selective axon regeneration. Top: In 5 dpf larval zebrafish (schematized in **A**), dorsal (green) and ventral (magenta) motor nerves exit the spinal cord in each body hemisegment via the ventral motor exit point (MEP) and diverge at a stereotyped branch-point to innervate the dorsal and ventral muscles, respectively. (**B**) Magnified schematic of single motor nerve in dashed box in (**A**) pre-transection, showing dorsal (green) and ventral (magenta) branches of the spinal motor nerve and associated Schwann cells (orange). Purple box, transection site. (**C**) At 8-15 hours post transection (hpt), growth cones enter the transection gap, and a small subset of Schwann cells at the nerve branch-point upregulate *col4a5* and *slit1a* (blue with red stripes). (**D**) By 48hpt, the majority of dorsal axons regenerate into the dorsal ROI, defined as 20-50° with respect to spinal cord. Bottom: Representative images of *Tg(isl1:GFP)*^+^ dorsal nerves at 5dpf and 48hpt, respectively in wildtype siblings (**E**, **F**) and *Tg(sox10:col4a5-Myc)* larvae (**H**,**I**).Yellow boxes, transection sites; green arrowheads, dorsal regrowth; magenta arrowheads, “errors” that regrew along non-dorsal paths. Scale bars, 10um. Sholl diagrams showing overlay of all *Tg(isl1:GFP)*^+^ fascicle trajectories at 48 hpt in wildtype siblings (**G**, n = 96 fascicles) and *Tg(sox10:col4a5-Myc)* larvae (**J**, n = 133 fascicles). For wildtype siblings, fascicles were counted for n = 37 nerves in 11 larvae; for *Tg(sox10:col4a5-Myc)* larvae, fascicles were counted for n = 40 nerves in 11 larvae. Green lines represent fascicles inside dorsal ROI; magenta lines represent fascicles outside of the dorsal ROI. Proportion of fascicles inside the dorsal ROI at 48 hpt was compared between *Tg(sox10:col4a5-Myc)* larvae and wildtype siblings using one-tailed Fischer’s exact test, (p = 0.0002). (K) Graph of regeneration error rate for wildtype siblings (No Tg) and *Tg(sox10:col4a5-Myc)* larvae (Tg^+^) compared using one-tailed t-test (p = 0.0001). Each dot represents error rate for one nerve; plus sign marks the mean; *** denotes p < 0.001.

To test this model, we first used genetic mutants of the Slit-Robo signaling pathway. We find that the Slit-receptor *roundabout2* (*robo2*) and the Slit-Robo co-receptor heparan sulfate (HS) are dispensable for target selection regeneration of ventral nerve axons, but are required to direct regenerating dorsal nerve axons at the branch choice-point. Moreover, we find that *robo2* is expressed in dorsal nerve neurons, that forcing robo2 expression in ventral nerve neurons is sufficient to redirect their regenerating axons into the dorsal nerve branch, and that this process requires Col4a5. Finally, using live cell imaging, we demonstrate that during regeneration *robo2* exerts its function at the nerve branch-point, preventing and correcting aberrant axonal extension of dorsal nerve axons, thereby promoting growth towards their original, dorsal targets. Combined our results reveal a previously unappreciated role for Slit-Robo signaling in axonal error prevention and correction, critical to ensure fidelity of branch selection and hence target selective regeneration.

## Results

### *Col4a5* upregulation at the nerve branch-point is critical to guide regenerating dorsal axons

We previously demonstrated that the glycosyltransferase *lysyl-hydrosylase-3* (*lh3*) and its substrate *collagen-4-alpha-5* (*col4a5*) are required to direct regenerating dorsal nerve axons towards their original targets ^12^. *Lh3* is constitutively expressed at low levels and acts in Schwann cells to promote target selective regeneration, while *col4a5* expression is transiently up-regulated 8-15 hours post transection (hpt) in a small subset of Schwann cells near the nerve branch-point (Fig. 1C and ref. ^12^, suggesting that *col4a5* expression restricted to the nerve branch-point might be instructive in directing regenerating axons. To test this idea, we generated a transgenic line, *Tg(sox10:col4a5-Myc)*, in which *col4a5* is now expressed in all Schwann cells, prior to and following peripheral nerve transection ^12^. Prior to nerve transection, dorsal nerves in wildtype siblings appear indistinguishable from those in transgenic animals expressing *col4a5* in all Schwann cells (compare Fig. 1E to H). Specifically, we quantified targeting of *Tg(isl1:GFP)*^+^ dorsal nerve axons prior to nerve transection in 5 day old animals. Dorsal nerve axons tightly fasciculate with one another, precluding us from quantifying individual axons contained in *Tg(isl1:GFP)*^+^ dorsal nerves. We therefore quantified the number of discernable *Tg(isl1:GFP)*^+^ fascicles and determined the fraction of fascicles within the dorsal muscle target area, which we previously defined as spanning 30° prior to transection (dorsal ROI Fig. 1D; ^12^). When we compared these fractions across genotypes, we found no significant difference in dorsal fascicles between *Tg(sox10:col4a5-Myc)* animals and wildtype siblings (Table S1), suggesting that in *Tg(sox10:col4a5-Myc)* animals, dorsal nerve targeting during development is unaffected.

To determine whether spatially restricted expression of *col4a5* is critical for target selective regeneration, we laser transected dorsal nerves in wildtype and *Tg(sox10:col4a5-Myc)* larvae and compared target selective regeneration at 48 hpt, when wildtype motor axons have re-established functional connections with their muscle targets ^20^. In wild type animals, 70% of fascicles containing dorsal nerve axons re-generated to their original dorsal target area (Fig. 1F, G; 37 nerves from 11 larvae), consistent with previous findings that regenerating axons readily select their original branch and targets ^12^. In contrast, transgenic expression of *col4a5* in all Schwann cells reduced target selective regeneration significantly. In *Tg(sox10:col4a5-Myc)*-expressing larvae, only 46% of the fascicles containing dorsal nerve axons selected their original dorsal trajectory (40 nerves from 11 larvae), concomitant with an 1.8 fold increase from 30% to 54% fascicles selecting incorrect ventral and lateral trajectories (Fig. 1I-J; Table S1; p = 0.0002, Fischer’s exact test). We next examined the regeneration error rate for individual nerves at 48 hpt and found that, when compared to wildtype siblings, nerves in *Tg(sox10:col4a5-Myc)*-expressing larvae displayed errors at a significantly higher rate (Fig. 1K; p = 0.0001, one-tailed t-test). Thus, expanding col4a5 expression from a small subset of Schwann cells strategically positioned at the nerve branch region to all Schwann cells impairs target selective regeneration. Moreover, the resulting phenotype closely mirrors the phenotype in mutants lacking *col4a5* ^12^. These results support the idea that *col4a5*’s transient expression in a subset of Schwann cells at the nerve branch point where regenerating axons of the dorsal branch select their branch specific trajectory is of functional importance.

### Slit-Robo signaling is required for target selective regeneration

The same small group of Schwann cells that up-regulates *col4a5* post injury concurrently upregulates slit1a expression ^12^, suggesting that similar to *col4a5*, *slit1a* might play a functional role in target selective regeneration. *Slit1a* encodes a canonical ligand for the Roundabout (Robo) family of repulsive axon guidance receptors ^26^, and is the only one of the four Slit ligands whose injury induced expression mirrors that of *col4a5* ^12^. Moreover, in the developing zebrafish visual system, Col4a5 directly binds to and is required for basement membrane anchoring of Slit, which guides laminar targeting of retinal ganglion cell axons through *robo2* ^25^. We therefore wondered whether *robo2* is involved in guiding regenerating dorsal axons. Using whole mount fluorescent *in situ* hybridization (ISH), we detected *robo2* mRNA expression in *Tg(isl1:GFP)*^+^ motor neurons of the dorsal nerve prior to transection and also during regeneration (Fig. S1).

We next asked whether Slit-Robo signaling plays a functional role in target selective regeneration. For this we examined dorsal nerve regeneration in genetic mutants for two Slit-Robo signaling components: mutants for *exotosin-like-3* (*extl3*), which lack Heparan sulfate (HS) ^27^, a glycosaminoglycan critical to stabilize Slit-Robo binding ^28^, and mutants for the Robo-receptor *roundabout2* (*robo2*). We first examined dorsal nerve regeneration in extl3 mutants, which at 5 dpf lack detectable levels of HS ^27^. Prior to nerve transection, targeting of dorsal nerve axons in 5 dpf day old *extl3* mutants was indistinguishable from their siblings (compare Fig. 2A to D; quantified in Table S1). In wildtype siblings, 68% of regenerating dorsal nerve fascicles returned to their original, dorsal targets (Fig. 2B-C), while 32% selected ventral and ventro-lateral trajectories (38 nerves from 19 larvae). In contrast, in *extl3* mutants we observed a 1.6 fold increase (from 32% to 52%) of dorsal nerve fascicles failing to select their dorsal trajectory, instead extending along erroneous ventral or ventro-lateral trajectories (23 nerves from 7 larvae)(Fig. 2E-F; Table S2, p = 0.0105, Fischer’s exact test). Similarly, when compared to wildtype siblings, individual nerves in extl3 mutants formed errors at a significantly higher rate at 48 hpt (Fig. 2G; p = 0.003, one-tailed t-test).

**Fig. 2.**
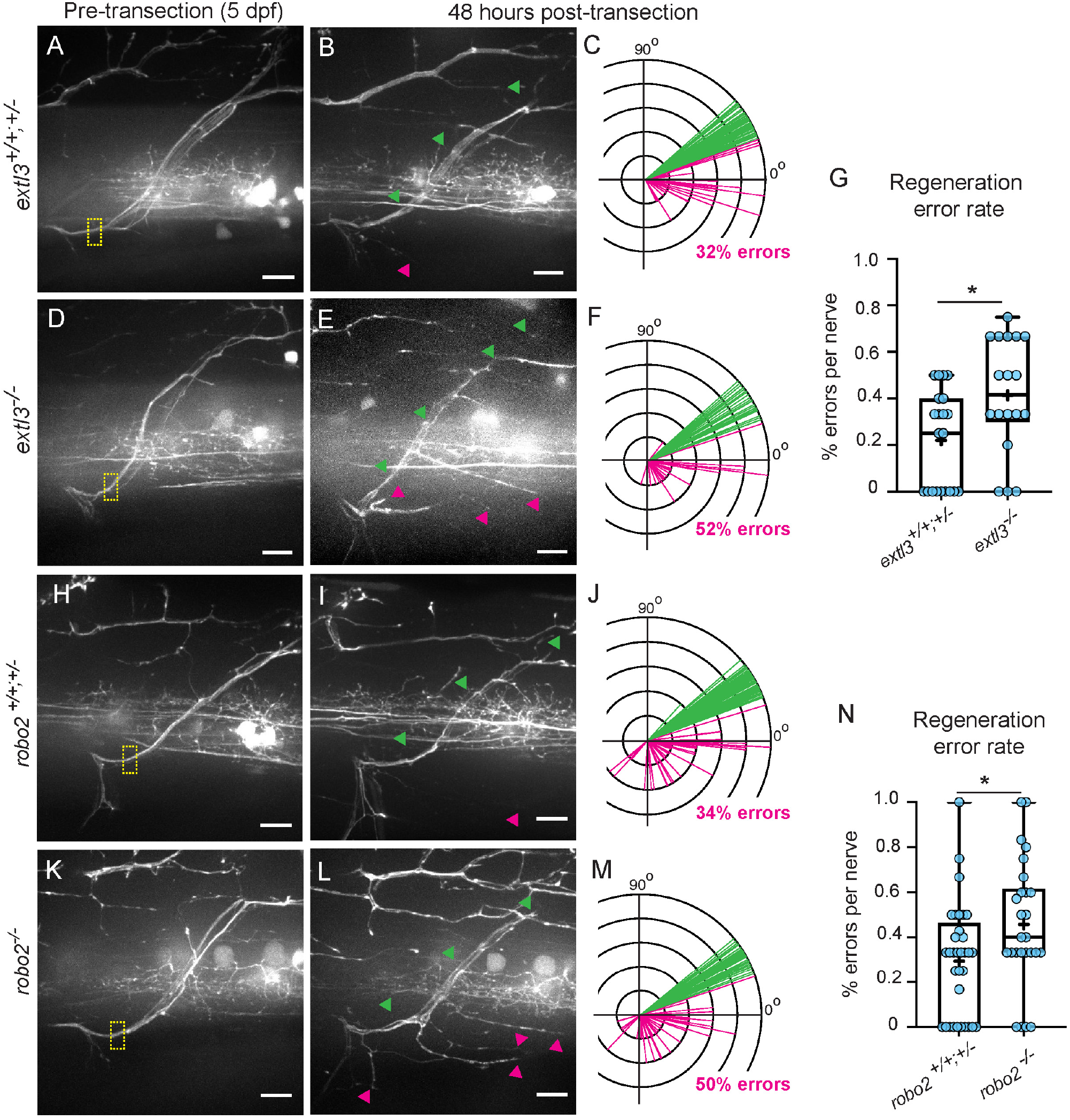
Slit-Robo signaling is required to guide regenerating dorsal nerve branch axons. Representative images of dorsal nerves in *extl3* wildtype siblings at 5 dpf and 48 hpt, respectively in *extl3* wildtype siblings (**A**, **B**), *extl3*^*−/−*^ (**D**, **E**), *robo2* wildtype siblings (**H**, **I**), *robo2*^*−/−*^ (**K**,**L**). Yellow boxes, transection sites; green arrowheads, dorsal regrowth; magenta arrowheads, “errors” that regrew along ventral and ventro-lateral paths. Scale bars, 10um. Sholl diagrams showing overlay of all *Tg(isl1:GFP)* fascicle trajectories at 48 hpt in *extl3* wildtype siblings (**C**, n = 98 fascicles), *extl3*^*−/−*^ (**F**, n = 58 fascicles), *robo2* wildtype siblings (**J**, n = 108 fascicles), *robo2*^*−/−*^ (**M**, n = 97 fascicles). Green lines represent fascicles inside dorsal ROI; magenta lines represent fascicles outside of the dorsal ROI. For *extl3* wildtype siblings, fascicles were counted in n = 38 nerves in 19 larvae; for *extl3*^*−/−*^ larvae, fascicles were counted in n = 23 nerves in 7 larvae; for *robo2* wildtype siblings, fascicles were counted in n = 33 nerves in 16 larvae; for *robo2*^*−/−*^ larvae, fascicles were counted in n = 26 nerves in 9 larvae. Proportion of fascicles inside the dorsal ROI at 48 hpt was compared using one-tailed Fischer’s exact test between *extl3* wildtype siblings and *extl3*^*−/−*^ larvae (p = 0.0105) and *robo2* wildtype siblings and *robo2*^*−/−*^ larvae (p = 0.0134). (G) Graph of regeneration error rate for *extl3* wildtype siblings and *extl3*^*−/−*^ larvae compared using one-tailed t-test (p = 0.003). (N) Graph of regeneration error rate for *robo2* wildtype siblings and *robo2*^*−/−*^ larvae compared using one-tailed t-test (p = 0.0116). Each dot represents error rate for one nerve. * denotes p < 0.05.

We next examined the role of *robo2* in dorsal nerve regeneration. Prior to transection, dorsal nerve targeting in *robo2* mutant animals was slightly lower than what we observed in robo2 wildtype siblings (compare Fig. 2H to K; quantified in Table S1), yet still within the range we have previously observed in wild type animals ^12^. Following nerve transection in wild type siblings, 34% of dorsal nerve fascicles failed to select their original trajectory, while the vast majority returned to their original target area (66%; Fig. 2I-J). In contrast, in *robo2* mutants only 50% of regenerating dorsal nerve fascicles returned to their original target area, an almost 1.5 fold increase of *Tg(isl1:GFP)*^+^ fascicles now extending along aberrant ventral or ventro-lateral trajectories (Fig. 2L-M; Table S2, p = 0.0134, Fischer’s exact test). Similarly, when compared to nerves in wildtype siblings at 48 hpt, individual nerves in *robo2* mutants exhibited significantly higher error rates (Fig. 2N; p = 0.0116, one-tailed t-test). Together these results demonstrate that Slit-Robo signaling plays a functional role in directing regenerating dorsal nerve axons along their original, pre-injury trajectories. Finally, to determine whether *extl3* and *robo2* play a selective role in promoting target selection of dorsal, rather than ventral nerve axons, we transected ventral nerves in *extl3* and *robo2* mutants. At 48hpt, ventral nerves in *extl3* and *robo2* mutants are indistinguishable from their siblings (Fig. S2, demonstrating that Slit-Robo signaling is selectively required for dorsal nerve target selective regeneration.

### *Robo2* promotes target selective regeneration at the nerve branch-point by preventing and correcting errors

To further understand the cellular mechanisms by which *robo2* promotes target selective regeneration, we examined the dynamics of regenerating axons navigating the nerve branch-point in *robo2* mutants. We have previously shown that after dorsal nerve transection in wildtype larvae, regenerating axons pause at the nerve branch-point and extend growth cones towards their original dorsal targets, as well as along erroneous ventral and lateral trajectories. Over the next few hours erroneous projections are destabilized, while growth cones along the dorsal path stabilize and continue to extend towards their original targets ^12^. To determine whether *robo2* directs regenerating dorsal nerve axons early in the process by minimizing the formation of erroneous projections, or subsequently by destabilizing already extending erroneous projections, we performed time-lapse imaging between 8 and 20 hpt as regenerating *robo2* mutant dorsal nerve axons navigate the branch choice-point (Figure 3, Movies S1 and S2). From these movies, we quantified the number of erroneous projections (errors), defined as *Tg(isl1:GFP)*^+^ growth ≥1μm that extended from the nerve branch-point along erroneous ventral or lateral trajectories (see Methods for more details). When compared to wild type siblings, *robo2* mutants exhibit no significant difference in the number of errors that form at the branch-point (Fig. S3). To determine whether there was a deficit in error correction at the branch-point in *robo2* mutants, we counted the number of errors (magenta arrowheads, Fig. 3, Movies S1 and S2) that were corrected. Errors were counted as “corrected” if they retracted within <1μm away from the nerve branch-point. When compared to siblings, *robo2* mutants displayed a significant decrease in the percent of errors that were corrected during early regeneration (Figure 3I).

**Fig. 3.**
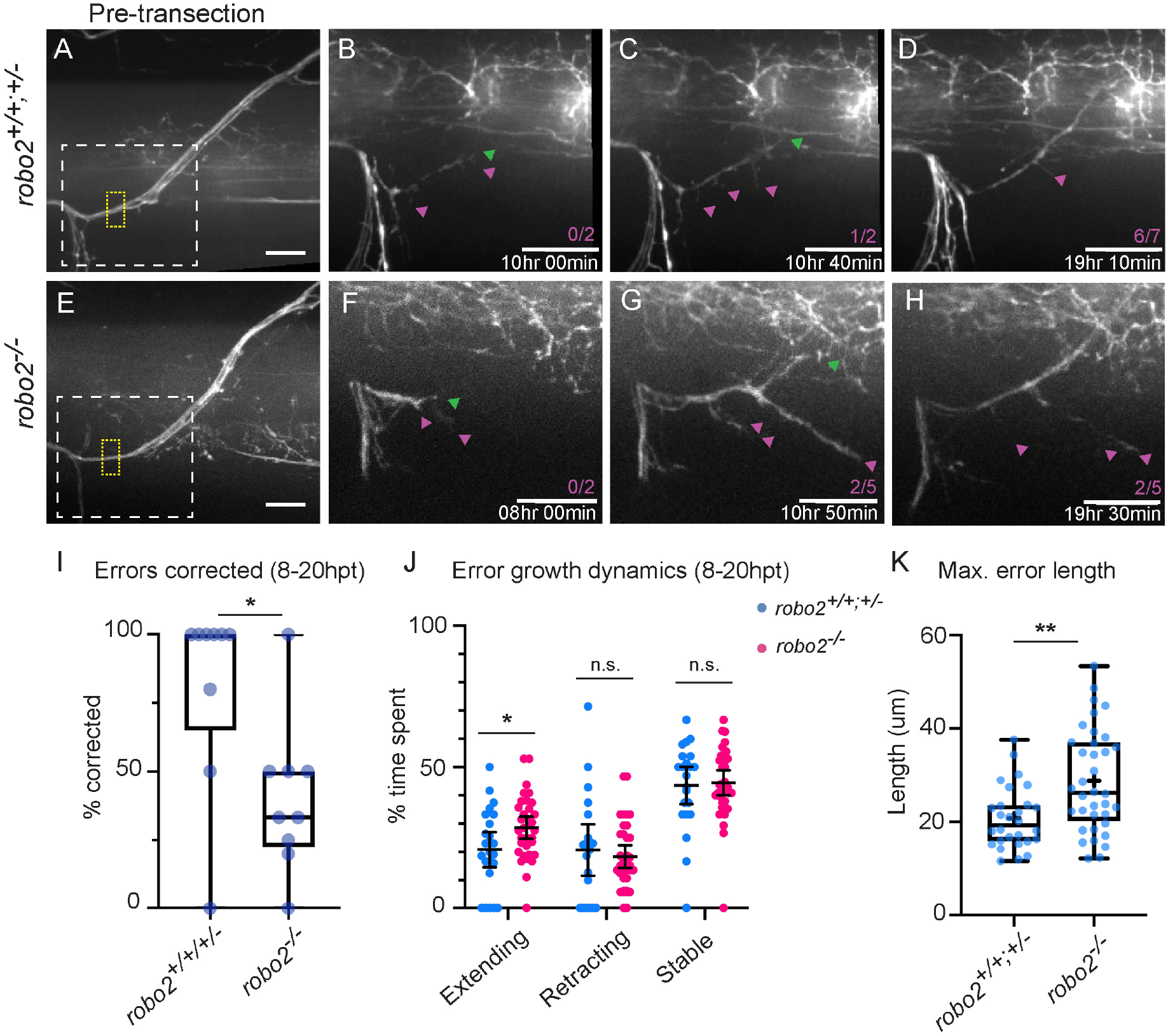
*robo2* prevents error extension at the nerve branch-point during regeneration. Representative images of *Tg(isl1:GFP)* nerves in *robo2*^+/−^ (wildtype sibling) (**A**) *robo2*^*−/−*^ (**E**) at 5dpf and during regeneration (**B-D**, **F-H**). Dashed yellow box, transection site; dashed white box, area enlarged 2x in B-D and F-H; green arrowheads mark dorsal regrowth; magenta arrowheads mark errors. Time post transection is denoted by white text in lower right of B-D and F-H. Errors are counted in magenta text in lower right of B-D and F-H as a fraction of [errors formed/errors corrected]. Scale bars, 10um. (I) Percent of errors corrected 8-20 hpt in wildtype siblings (*robo2*^+/+^;^+/−^, n = 9 nerves) and *robo2*^*−/−*^ larvae (n = 10 nerves). Errors were counted as “corrected” when their length measured from the nerve branch-point was <1um. Each dot represents one nerve. Ranks were compared between genotypes using two-tailed Mann Whitney test (p = 0.0208). (J) Quantification of regenerating axon dynamics in wildtype siblings and *robo2*^*−/−*^ larvae 8-20 hpt plotted by total time spent extending, retracting, and stable (no movement). Each dot represents one error (for siblings, n = 23 errors; for *robo2*^*−/−*^, n = 34 errors). Error movements were examined in 10 min intervals and classified as extensions when there was a 1um increase in error length measured from the motor exit point (MEP); movements were classified as retractions when ≥1um decrease in error length measured from the MEP; errors were classified as stable when no movement ≥1um occurred. Line, mean; error bars, 95% confidence intervals. Means were compared between genotypes using two-tailed t-tests (extending, p = 0.0365; retracting p = 0.9131; stable, p = 0.2491). (**K**) Maximum length of errors in wildtype siblings and *robo2*^*−/−*^ larvae in ums measured from the motor exit point (MEP). Each dot represents one error. Plus sign marks the mean. Means were compared between genotypes using two-tailed t-test (p = 0.0012). ** denotes p < 0.01; * denotes p < 0.05; n.s. denotes “not significant.”

Based upon the well characterized role of Slit-Robo in axon repulsion ^26^, we considered whether in *robo2* mutants the deficit in error correction at the nerve branch-point might be due to reduced error retraction. To test this, we measured in wildtype siblings and *robo2* mutants the percent of time that errors spent retracting, extending or being stable (no movement). We failed to detect any significant difference in relative time that errors spent retracting in *robo2* mutants compared to wild type siblings (Fig. 3J). Similarly, we did not observe any differences in the average speed of extension or retraction in *robo2* mutants when compared to siblings (Fig. S3B). Instead, compared to wildtype siblings, erroneous projections (errors) in *robo2* mutants spent significantly more time extending (Fig. 3J). This is the result of a combined reduction in the time that errors spent retracting and stable in *robo2* mutants, neither of which is statistically significant on its own (Fig. 3J). These results suggest that rather than promoting axonal retraction along incorrect trajectories, *robo2* promotes dorsal nerve target selection by preventing axon extension along erroneous ventral and lateral trajectories. Consistent with this idea, when compared to wild type siblings, erroneous projections in *robo2* mutants grew longer distances (Fig. 3K). Combined with previous results this provides strong support for a model in which *robo2* expression on regenerating dorsal prevents and corrects errors at the nerve branch-point in response to Slit1a transiently produced by a small subset of adjacent Schwann cells and spatially scaffolded by Col4a5. By preventing error extensions, *robo2* tilts the balance between extension and retraction such that errors retract more often than they extend. This results in shorter errors that are readily corrected through *robo2*-independent mechanisms of retraction, ultimately biasing regenerating dorsal nerve axons towards their original dorsal trajectory.

### *Robo2* expression drives target selective regeneration

We next asked how Slit-Robo signaling selectively influences regeneration of dorsal but not ventral nerve axons. One possibility is that *robo2* functions selectively in dorsal nerve axons, enabling regenerating axons of only the dorsal but not the ventral branch to mount a *slit1a*-dependent error response at the nerve branch-point. We hypothesized that if this were the case, forcing *robo2* expression in re-generating ventral nerve axons would redirect them onto a dorsal trajectory. To test if *robo2* expression is indeed sufficient to drive target selective regeneration, we used the motor neuron-specific mnx1 promotor ^29^ to transiently express mKate alone or mKate with *robo2* in small subsets of motor neurons (for more details see Methods). Importantly, when compared to mKate expression, *robo2*-mKate expression in motor neurons did not impair their ability to grow nor did it change their developmental bias in selecting a ventral or dorsal trajectory (Fig. S4). This is consistent with the absence of a developmental motor axon phenotype in robo2 mutants (Table S1), further confirming that *robo2* acts selectivity during the regeneration process. To determine whether *robo2* is sufficient to promote dorsal branch selection in regenerating axons, we screened for spinal motor nerves with small subsets of mKate^+^ axons along the ventral, but not the dorsal branch (Fig. 4A and 4C). We laser transected these ventral nerves and assessed the regeneration of mKate^+^ fascicles at 48 hpt. We found that regenerating mKate^+^ ventral nerve axons always selected a ventral path towards their original targets (Fig. 5B; Table S3, n = 13/13 nerves), consistent with previous results ^20^. In contrast, forcing *robo2*-mKate expression in regenerating ventral nerve axons was sufficient to redirect them onto a dorsal trajectory (Fig. 4C-D; Table S3, n = 7/15 nerves, Fischer’s exact test p=0.0069). Importantly, the trajectories taken by these axons were indistinguishable from those taken endogenously by dorsal nerve axons (compare to Fig. 2B, I). Thus, *robo2* is both required and sufficient to drive target selective regeneration.

**Fig. 4.**
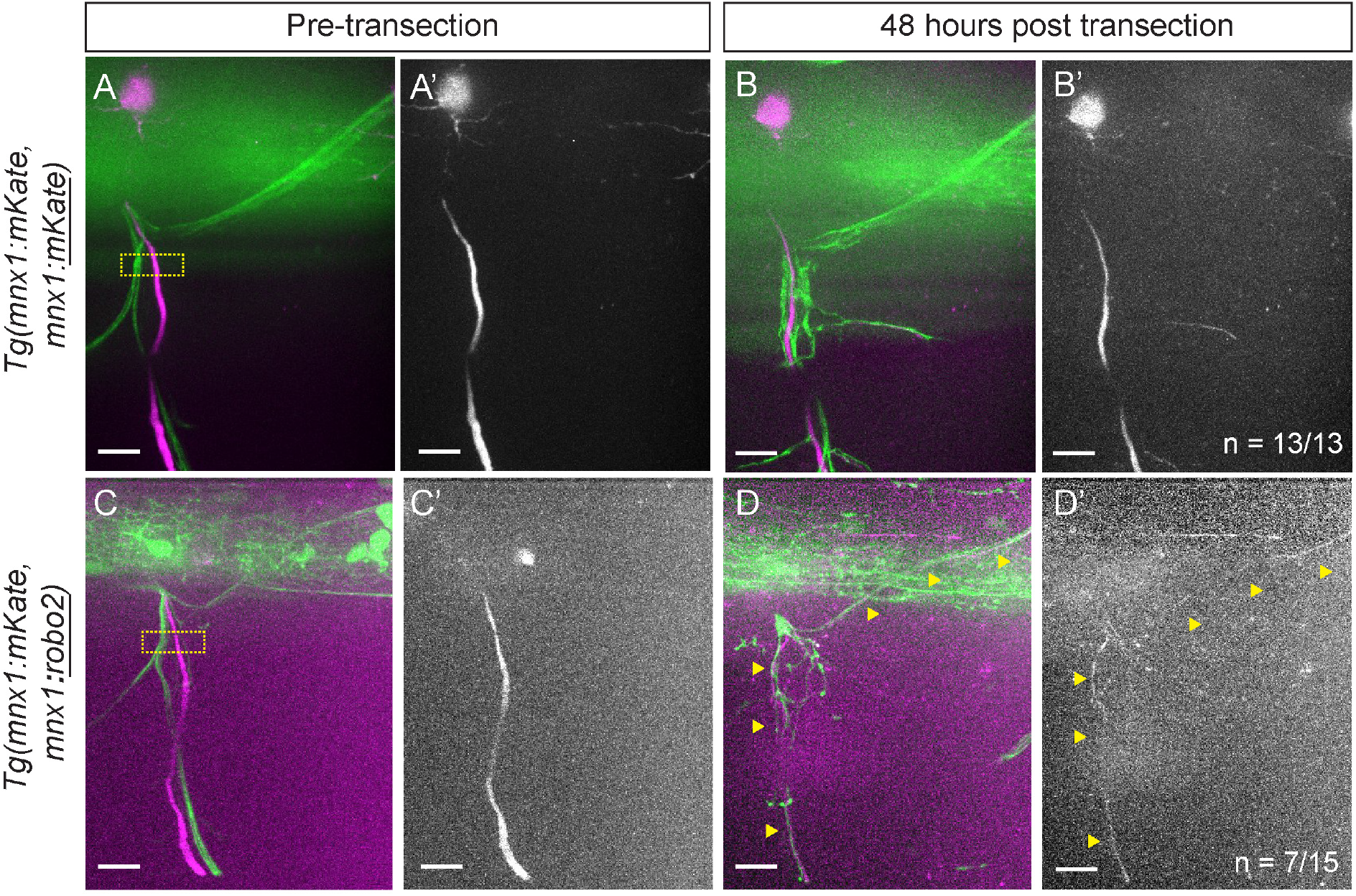
*robo2* is sufficient to promote dorsal branch-selection by regenerating axons. Representative images of *Tg(hb9:GFP)* (green) nerves with small numbers of fascicles expressing transient *Tg(mnx1:mKate, mnx1:mKate)* (**A-B**) or *Tg(mnx1:mKate, mnx1:robo2)* (**C-D**) (magenta) in small subsets of ventrally projecting motor axons. Merged GFP and mKate images shown at 5dpf (A, C) and 48hpt (B, D). mKate channel is shown alone at 5dpf (**A’**, **C’**) and 48hpt (**B’**, **D’**). In larvae expressing *Tg(mnx1:mKate, mnx1:mKate)*, n = 13/13 nerves had only ventral regrowth of mKate^+^ fascicles, like in the example shown. In larvae expressing *Tg(mnx1:mKate, mnx1:robo2)*, n = 7/15 nerves had dorsal regrowth of mKate^+^ fascicles, like in the example shown. Proportions of dorsal regrowth were compared between conditions using one-tailed Fischer’s exact test (p = 0.0054). Images were processed as described in Materials and Methods. Dashed yellow boxes, transection site. Scale bars, 10um.

**Fig. 5.**
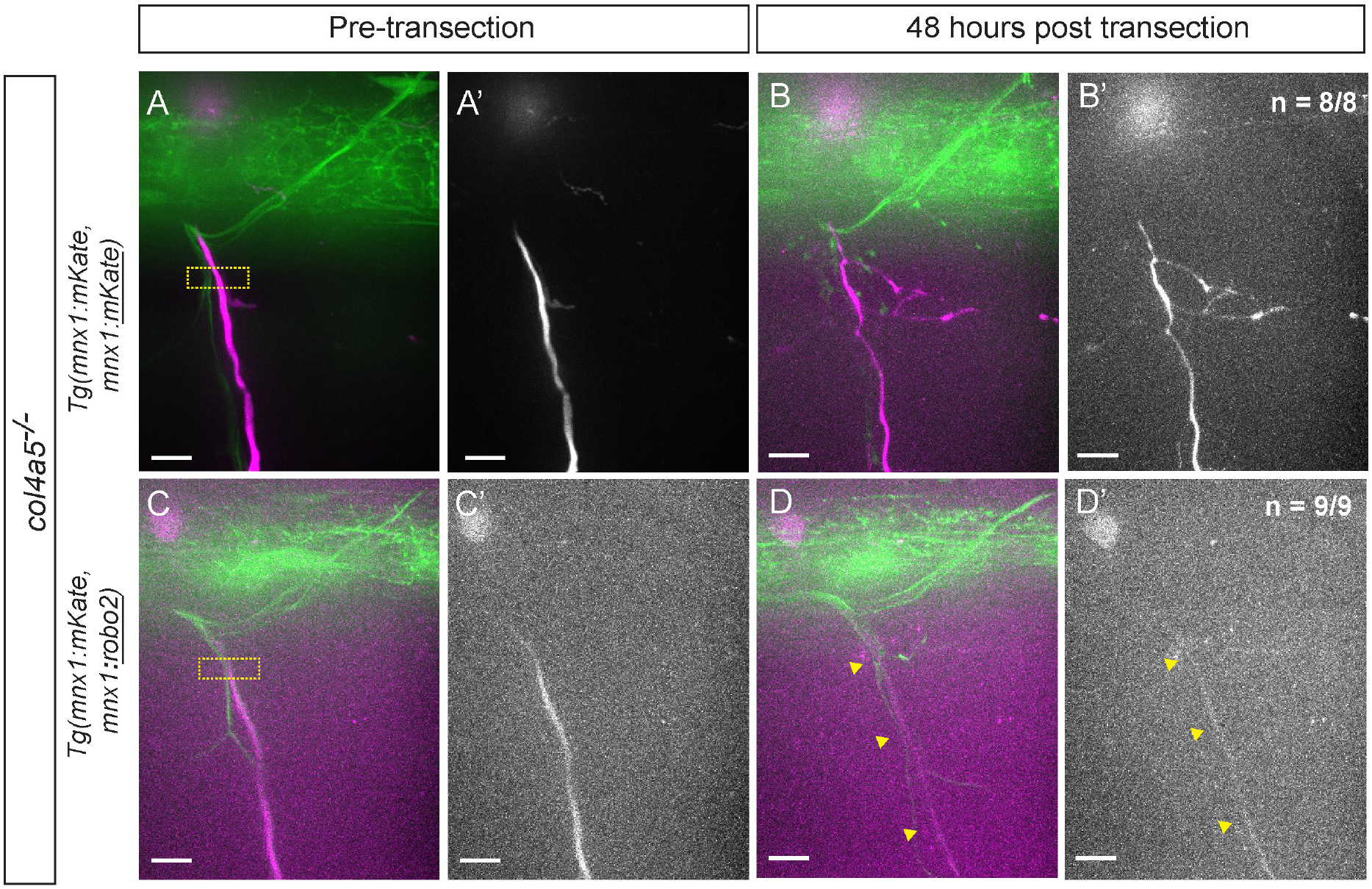
*col4a5* is required for the role of *robo2* in branch-selective axon regeneration. Representative images of *Tg(hb9:GFP)* (green) nerves in *col4a5*^*−/−*^ larvae with fascicles expressing transient *Tg(mnx1:mKate, mnx1:mKate)* (**A-B**) or *Tg(mnx1:mKate, mnx1:robo2)* (**C-D**) (magenta) in small subsets of ventrally projecting motor axons. Merged GFP and mKate images shown at 5dpf (A, C) and 48hpt (B, D). mKate channel is shown alone at 5dpf (**A’**, **C’**) and 48hpt (**B’**, **D’**). In larvae expressing *Tg(mnx1:mKate, mnx1:mKate)*, n = 8/8 nerves had only ventral regrowth of mKate^+^ fascicles, like in the example shown. In larvae expressing *Tg(mnx1:mKate, mnx1:robo2)*, n = 9/9 nerves had only ventral regrowth of mKate^+^ fascicles, like in the example shown. Proportions of ventral regrowth were compared between conditions using one-tailed Fischer’s exact test (p = 0.999). Images were processed as described in Materials and Methods. Dashed yellow boxes, transection site. Scale bars, 10um.

### *Robo2* requires *col4a5* function for target selective regeneration

In *robo2* and *col4a5* mutants, ventral branch axons reliably regenerate along their appropriate ventral path (Fig. 4; ref. ^12^), while dorsal branch axons frequently fail to select their original dorsal trajectory and instead extend along erroneous, ventral and lateral trajectories. Because of the similarities of their mutant phenotypes, we next asked whether *robo2* and *col4a5* act through two distinct pathways or whether they are both part of one common pathway. We reasoned that if the latter was the case, then redirecting ventral nerve axons towards dorsal trajectories via forced *robo2* expression should depend on *col4a5* function. To test this hypothesis, we repeated the *robo2* mis-expression experiment driving sparse expression of either mKate or *robo2*-mKate in small subsets of regenerating ventral nerve axons, but now in a *col4a5* mutant background. Prior to nerve transection at 5 dpf, there was no difference between the branch-selection of sparsely labeled wildtype and *robo2*-expressing axons in *col4a5* siblings or mutants (Fig. S5). This is consistent with our previous findings that *col4a5* is dispensable for spinal motor nerve development ^12^. Like before, we selected spinal motor nerves with small subsets of mKate^+^ axons along the ventral, but not the dorsal nerve branch, laser transected these ventral nerves and at 48 hpt assessed the regeneration of mKate^+^ axons. In *col4a5* siblings and mutants, regenerating ventral nerve axons expressing mKate faithfully selected their original, ventral trajectory (Fig. 5A-B; Fig. S6A-B; Table S3, *col4a5* siblings: n = 12/12; *col4a5* mutants n = 8/8). In *col4a5* siblings, *robo2*-mKate expression was sufficient to redirect regenerating ventral nerve axons onto a dorsal trajectory (Fig. S6C-D; Table S3, n = 5/14 nerves, Fischer’s exact test p=0.0425). In contrast, in *col4a5* mutants, *robo2*-mKate expression failed to redirect ventral nerve axons onto a dorsal trajectory (Fig. 5C-D; Table S3, n = 0/9), demonstrating that *col4a5* function is required for *robo2* to redirect ventral nerve axons dorsally. This provides compelling evidence that *col4a5* and Slit-Robo act in a common genetic pathway that promotes dorsal branch selection of regenerating axons. Combined, our results support a model in which a small subset of Schwann cells strategically located at the branch choice point upregulate the expression of *col4a5* and *slit1a* in response to nerve injury. As regenerating axons approach the branch choice point, Robo2 function selectively in dorsal nerve axons prevents and correct erroneous projections along ventral and lateral trajectories, thereby biasing axonal regrowth towards their original, dorsal trajectory (Fig. 6).

**Fig. 6.**
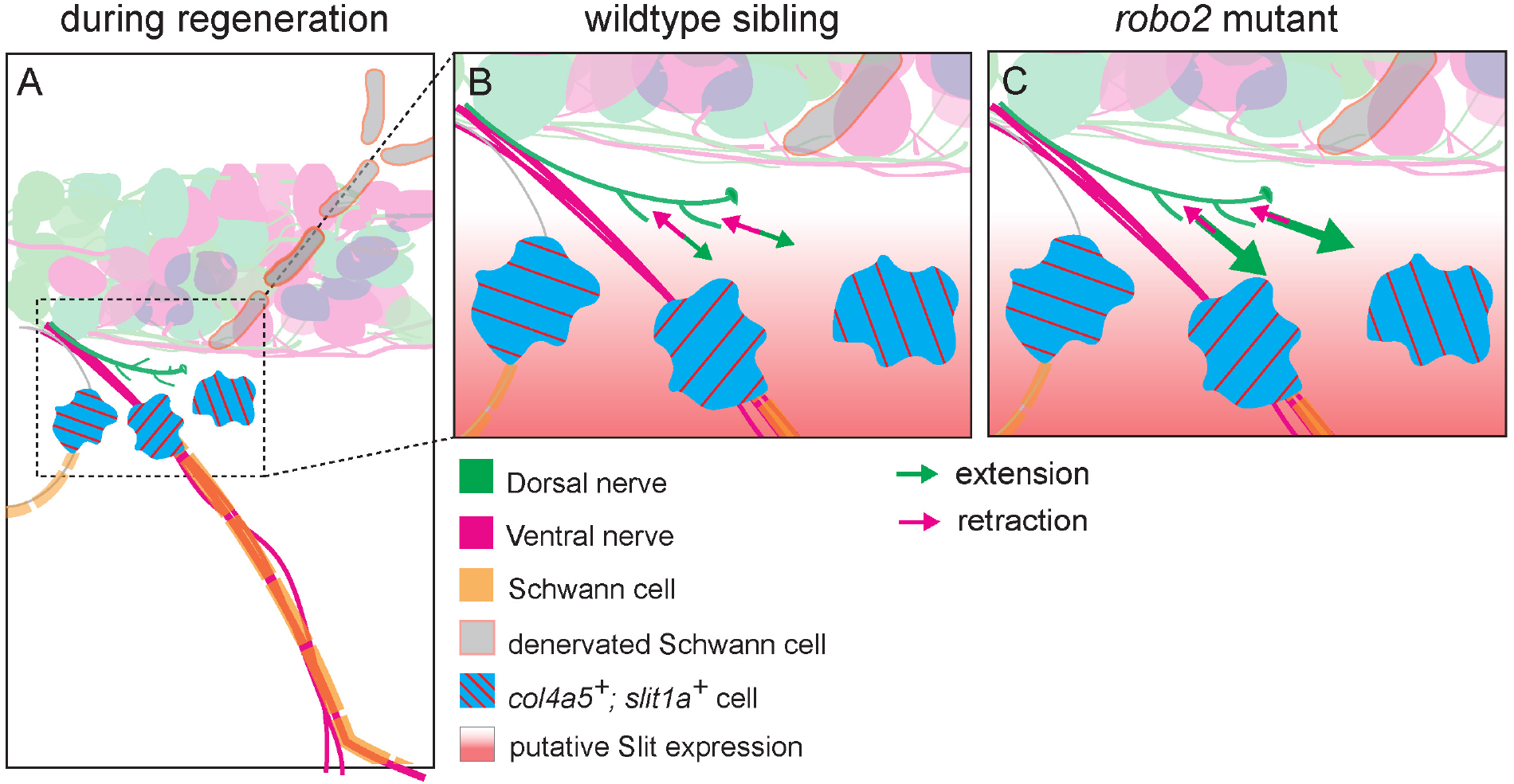
Model for role of *col4a5-robo2* pathway in target-selective axon regeneration. During regeneration, (**A**) a subset of specialized Schwann cells at the nerve branch-point express *col4a5* (blue with red stripes), which scaffolds the repulsive axon guidance cue Slit (red gradient) in the local ECM. (**B**) In response to Slit at the nerve branch-point, regenerating wildtype axons expressing the Slit-receptor robo2 extend for short distances along ventral and ventro-lateral paths, as they navigate the nerve branch choice-point. These short erroneous extensions (green arrows) are balanced by retraction events (red arrows), which results in their eventual retraction and correction. (**C**) in *robo2*^*−/−*^ larvae, regenerating axons are not responsive to Slit at the nerve branch-point and thus extend more frequently along ventral and ventro-lateral paths. These frequent erroneous extensions (large green arrows) are not balanced by retraction events (red arrows), which occur with similar frequency as retractions in wildtype siblings. Thus, in *robo2*^*−/−*^ larvae, branch-point errors extend for longer distances and are less likely to be corrected.

## Discussion

In response to peripheral nerve injury, regenerating axons face the challenge of navigating toward and reconnecting with their original synaptic targets. The difficulty of this task increases with the architectural complexity of the injured nerve. After exiting the spinal cord, peripheral nerves repeatedly divide into progressively smaller branches, frequently leading to different targets ^11^. Regenerating axons may therefore encounter multiple nerve branch-points where they confront the choice to select their appropriate, preinjury trajectory. This task is further compounded by an environment that is dramatically different from the one they successfully navigated during development. Despite constituting an enormous navigational challenge, regenerating axons are able to preferentially select their original nerve branch ^12,15,30^, and to regrow toward appropriate targets ^31–34^. Combined, previous studies strongly support the notion that regenerating axons are guided at nerve branch-points by dedicated molecular mechanisms, yet few such mechanisms have been described. Here we identify a molecular pathway critical for communication between glia cells located at a nerve branch-point and regenerating axons to direct axons of one nerve branch onto their preinjury trajectory. Specifically, our results provide compelling evidence for transient, spatially restricted, and tightly coordinated signaling events between *col4a5/slit1a*-expressing Schwann cells and *robo2*-expressing regenerating axons at the nerve branch-point critical to promote target selective regeneration.

### *Robo2* selectively destabilizes erroneous axonal extension at the nerve branch-point

Live cell imaging experiments provide compelling evidence for a Robo2-dependent mechanism that directs regenerating axons into the appropriate nerve branch as they encounter the branch choice-point. As they encounter the nerve branch-point, regenerating dorsal nerve axons in both wildtype and *robo2* mutants initiate growth (~1μm) along erroneous ventral or ventro-lateral trajectories with similar frequencies (Fig. S3A). In contrast to wildtype, *robo2* mutant axons along these erroneous trajectories are significantly more likely to extend (Fig. 3J), ultimately growing for distances 1.5 to 3 times longer than those in the wildtype (Fig. 3K). Thus, rather than preventing initial error formation per se, *robo2* destabilizes erroneous axons (>~1μm), thereby preventing their further growth (Fig. 3K). Thus, while *robo2* destabilizes erroneous axonal growth, it promotes rather than inhibits regeneration. This is markedly different from the role that canonically repulsive axon guidance systems, including Slit-Robo, often play in inhibiting axon extension after injury, resulting in poor or stalled regeneration ^10,35^. For example, in C. elegans, Slit (*slt-1*) and Robo (*sax-3*) inhibit the extension of the mechanosensory PLM axon after transection, ultimately leading to reduced regeneration ^36^. Rather than negatively impacting axon regeneration, we find that growth rates of regenerating axon outgrowth in *robo2* mutants are identical to those in wild type animals (Fig. S3B). This provides strong evidence that for zebrafish peripheral nerve regeneration *robo2*-independent mechanisms promote axon outgrowth, while *robo2*’s role is to selectively bias regenerative growth of dorsal nerve axons toward their original trajectory.

How similar is this *robo2*-dependent mechanism to other, well documented mechanisms known to promote target selective regeneration in mammals? It is well documented that in mammals motor axons preferentially regenerate into their original nerve branches ^13–15^, and that this process is regulated by Schwann cells and by nerve end organs such as muscle and skin ^37,38^. For example, after injury, Schwann cells in the distal stump upregulate branch-specific neurotrophic factors and cell adhesion molecules ^39–42^. These molecules support the outgrowth and maintenance of appropriate axonal populations ^43,44^ such that when axons regenerate into inappropriate nerve branches, the resulting errors are pruned away over weeks or months ^30,45^. Thus, in contrast to well documented pruning mechanisms occurring long after axons have regenerated towards incorrect targets, the *robo2*-dependent mechanism we describe here is engaged during the time period when regenerating axons are confronted at a choice point with the task to select their original trajectory, thereby promoting target selective regeneration locally and on a much shorter timescale.

### A Co4a5/glia Robo2/axon dependent mechanism provides local guidance to promote target selective regeneration

In response to peripheral nerve injury, Schwann cells distal to the injury site dedifferentiate through a well characterized molecular pathway ^46,47^. These dedifferentiated Schwann cells exhibit differences in gene expression ^39–41^, yet the functional significance of these gene expression differences, including their functional roles in branch selection and thus target selective regeneration have remained largely unknown. We previously reported that in response to peripheral nerve injury in zebrafish, a small, selective group of Schwann cells (~1-3) strategically located at where dorsal nerve branch deviates from the ventral nerve branch, upregulate both col4a5 and its binding partner, the repulsive guidance cue *Slit1a* ^12,25^. While we had shown that *col4a5* is critical for target selective regeneration, whether the post injury expression of *col4a5* spatially restricted to just a few Schwann cells was important for target selectivity was unclear. Similarly, whether Slit1a played a functional role in this process and whether Slit1a and *col4a5* expression were functionally related had not been defined.

Our results demonstrate that expanding the expression of *col4a5* to all Schwann cells severely impacts target selective regeneration (Fig. 1). Importantly, rather than extending along random trajectories, regenerating dorsal nerve axons now extended along erroneous ventral and ventro-lateral trajectory similar to those we observe in mutants lacking *col4a5*, *robo2* and *extl3*, respectively (Fig. 2; ref. ^12^). Moreover, in animals expressing *col4a5* in all Schwann cells, regenerating dorsal nerve axons made errors precisely where they pause and explore the nerve branch-point before turning dorsally during regeneration (Fig. 3). Combined, this provides compelling evidence that rather than providing a permissive substrate and environment, spatially restricted *col4a5* expression is critical to instruct regenerating dorsal nerve axons at the nerve branch-point.

Our results also reveal a previously uncharacterized role for role Slit-Robo signaling in target selective regeneration (Fig 5). Loss of function mutations in two Slit-Robo signaling components, *extl3* and *robo2*, result in the same target selective regeneration defects (Fig. 2). Conversely, we find that transgenic expression of *robo2* in ventral nerve axons, which are unaffected by the loss of Slit-Robo signaling, is sufficient to redirect these axons onto a dorsal nerve branch specific route (Fig. 4), and that this process requires Col4a5 function (Fig. 5). Given the incomplete penetrance of the *robo2* mutant phenotype in target selective regeneration, we cannot exclude the possibility that one or multiple of the additional three zebrafish Robo receptors ^48,49^, play a role in this process and hence partially compensate for the loss of *robo2*. Future experiments, including generating single and double mutant combinations for the other three Robo receptors as well for each of the four known Slit ligands ^50^ will provide a comprehensive view on role of Slit-Robo signaling during target selective regeneration and inform the contribution of individual Robo receptors in this process. Nonetheless, our results provide strong support for a mechanism in which glial-derived *col4a5* expression restricted to the branch-point promotes dorsal turning of regenerating axons possibly via Col4a5-bound Slit1a. Moreover, the transient expression of *col4a5* and *slit1a*, which only lasts a few hours and coincides with the time period when regenerating axons navigate the branch choice point is remarkable, underscoring the high degree and functional importance of spatially and temporally signaling between regenerating axons and Schwann cells to achieve target selective regeneration. While our results identify a previously uncharacterized molecular mechanism to promote peripheral nerve regeneration, they also draw attention to the need to incorporate spatially and temporally restricted deliveries of guidance information in therapeutic strategies aimed at enhancing target selective regeneration.

## Materials and Methods

### Fish Lines and Maintenance

All fish lines were maintained in Tügbingen or Tupfel longfin (TLF) backgrounds and maintained as previously described ^51^. We used the following mutant alleles, which were genotyped as previously described: *robo2-ti272z* ^52^, *extl3-tm70g* ^27^, *col4a5-s510* ^53^.The *Tg(sox10:col4a5-Myc)* line was generated as previously described ^12^ and genotyped by amplifying the Myc transgene using the following primers: 5’ GAC-TACAAGGATGACGATGACAAG 3’ (forward) and 5’ TTCTCCCATAGTCACGCTAGC 3’ (reverse). For *in vivo* imaging, the following transgenic lines were used: *Tg(mnx1:GFP)*^*ml*3 29^ to visualize motor axons in both dorsal and ventral nerve branches, *Tg(isl1:GFP)*^*rw*0 54^ to visualize dorsal nerve branch axons alone. Zebrafish veterinary care was performed under the supervision of the University Laboratory Animal Resources (ULAR) at the University of Pennsylvania. All zebrafish work was performed in accordance with protocols approved by the University of Pennsylvania Institutional Animal Care and Use (IACUC).

### Nerve Transection

Dorsal and ventral nerves were transected using a nitrogen-pulsed dye (440nm) laser as previously described ^20^. Briefly, one of the two nerve branches (dorsal for Figures 1, 2 and 3 or ventral for Figures 4 and 5) were transected ~5um from the nerve branch-point (10-15um from the MEP), leaving the other nerve branch intact and a ~5um gap between proximal and distal stumps of the transected nerve branch.

In *extl3*^−/−^ larvae, dorsal nerves reach dorsal muscle targets, but a small subset grow along an aberrantly lateral trajectory. This phenotype is variably penetrant, affecting 0-50% of nerves per larvae and <20% of all nerves across the mutant population (n>50 larvae; PLM, unpublished observations). Thus, for dorsal nerve transections in *extl3*^−/−^ larvae, we carefully selected phenotypically normal nerves for transection.

### Quantification of Axon Regeneration

Dorsal axon guidance pre- and post-transection was quantified using modified Sholl analysis, as previously described ^12^, with the exception that line thickness in Sholl diagrams here do not correlate to fascicle thickness. To calculate the regeneration error rate for each nerve, we divided the number of fascicles that regenerated outside the dorsal ROI (“errors”) by the total number of fascicles that regenerated by 48 hpt. To determine the angle of ventral nerve extension, we measured the angle between two consistent points along the trunk of the nerve (50um and 100um from the MEP). The extent of ventral nerve regeneration was scored by counting the number of fascicles which regenerated at least 50um from the MEP by 48 hpt.

### Immunohistochemistry and whole-mount fluorescent *in situ* hybridization

To visualize *robo2* expression after nerve transection, dorsal nerves were transected in 5 dpf *Tg(isl1:GFP)* larvae. At 0-10 hpt, larvae were fixed in 4% PFA in PBS with 0.1% Tween-20 overnight at 4°C. The antisense *robo2* probe was synthesized from pBlueScript-robo2 linearized with EcoRI using T3 RNA polymerase (Promega); the sense *robo2* probe was synthesized from pBlueScript-robo2 linearized with XhoI using T7 RNA polymerase (Promega). Probes were hydrolyzed in 0.6M sodium carbonate and 0.4M sodium bicarbonate at 60°C for 11min to yield 300-500bp fragments. Whole mount in situ hybridization was performed as previously described ^55^ with the following modifications: 5 dpf larvae were permeabilized by digesting with Proteinase K (10ug/ml, Promega) for 2 hours; endogenous peroxidases were quenched by incubating in 0.3% H_2_O_2_ for 30 minutes before adding anti-digoxigenin antibody; for blocking and antibody incubation, we used 2% Blocking Reagent (Roche) in PBS with 0.1% Tween-20; probes were detected using sheep anti-digoxigenin POD Fab fragments (Roche) and developed for 2 minutes using Tyramide Signal Amplification (TSA Plus kit, PerkinElmer). We stained for emphTg(isl1:GFP) using chicken anti-GFP (1:500, Aves Labs) detected by donkey anti-chick Alexa Fluor 488 (1:500, Jackson ImmunoResearch Laboratories, Inc.). Anti-digoxigenin POD and anti-GFP primary antibodies were incubated concurrently; secondary antibody was incubated at 4°C overnight after TSA.

Nerves were imaged in 1um sections using a 63X water immersion lens on a Zeiss LSM 880 laser scanning confocal microscope and Zeiss Zen software or using a 40X water immersion lens on an Olympus Spinning disk confocal microscope and 3i Slidebook Software. Overlap between emphTg(isl1:GFP) and emphrobo2 probe was quantified from 40X images in Fiji in the following way: motor pools of transected nerves were isolated by cropping optical sections in 3D; motor pools were then compressed into MIPs; GFP and robo2 probe signals were separated and converted into binary masks using max entropy and moments methods, respectively; percent particle overlap between the two masks was calculated using the GDSC colocalization plugin; percent particle overlap was then normalized to the number of cell bodies manually counted in each motor pool.

### Plasmid construction

The mnx1:mKate, mnx1:robo2 plasmid was constructed using Gateway cloning ^56^ using pME:robo2^57^ and pI-SceI mnx1:mKate, mnx1:DEST, which encodes two mnx1 promoters in tandem ^58^, to insert the *robo2* coding sequence behind the second mnx1 promoter. To construct plasmid for synthesis of *robo2 in situ* probe (pBlueScript-robo2), the full length *robo2* coding sequence was amplified from the mnx1:mKate, mnx1:robo2 plasmid using the following primers 5’ AGTCAGCTCGAGAACGTGTTCTGGGGT-TGAGA 3’ (forward, includes XhoI restriction site) and 5’GCTAAC-GAATTCTGGGTATGAGGCATTTCCAGAAC 3’ (reverse, includes EcoRI restriction site). XhoI and EcoRI restriction sites were used to clone this product into pBluescript II KS+.

### Sparse axonal labeling

We used mnx1:mKate, mnx1:robo2 and mnx1:mKate, mnx1:mKate to label small numbers of ventral motor neurons by injecting 50-100 pg of plasmid DNA into one-cell stage embryos with Isce-I, as previously described ^59^. We have previously validated that both mnx1 promoters in this construct are active and drive comparable levels of expression (ref. ^58^ and unpublished observations). At 3dpf, injected larvae with mKate expression were screened for ventral nerves with very few labeled axons using a 40X water immersion lens on a Olympus spinning disk confocal microscope using 3i Slidebook software. When examining developmental labeling frequency, nerves were scored as “dorsal” if any mKate^+^ fascicles were present were present along the dorsal branch, regardless of whether mKate^+^ fascicles were also present along the ventral branch.

It was technically challenging to sparsely label ventral axons in sufficient numbers for transection experiments without labeling multiple neurons per motor pool. We could almost always count multiple (2-6) mKate^+^ cell bodies in motor pools corresponding to transected ventral nerves (data not shown). After ventral nerve transection, we often observed that emphTg(mnx1:mKate;mnx1:robo2)^+^ fascicles regenerated along both the dorsal and the ventral branch (see Figure 5D). We believe it is very likely that this reflects the regeneration of multiple labeled axons, rather than single bifurcating axons. Therefore, at 48 hpt, we scored mKate^+^ fascicle regeneration as “ventral,” if mKate^+^ fascicles were observed only on the ventral nerve branch, and we scored mKate^+^ fascicle regeneration as “dorsal” if we observed any mKate^+^ along the dorsal branch, regardless of whether mKate^+^ fascicles were also present along the ventral branch.

### Live Imaging

Larvae were anesthetized, mounted in agarose and imaged on a spinning disk confocal microscope as previously described ^20^. We began our timelapse experiments 7-9 hpt and filmed regeneration for 12-15 hours. Due to variability in time when axons started regrowing (8-14 hpt), we quantified axon dynamics starting when the first regenerating fascicle reached the nerve branch-point and ending up to 10 hours later. We analyzed regenerating fascicles for a total of 5940 minutes in siblings (n = 9 nerves) and 5030 minutes in mutants (n = 10 nerves).

### Image Processing

For ventral nerves (Fig. S2) and fixed samples (Fig. S1), Z-stacks were compressed into maximum intensity projections (MIPs). Brightness and contrast were automatically optimized based upon the image histogram in Fiji ImageJ (NIH). The dorsal nerve branch wraps around the spinal cord, closely apposed to motor neuron cell bodies, which are labeled brightly by our transgenic lines. To visualize the dorsal nerve independently of neuron cell bodies (Figs. 1, 2, 3, 4, 5), we used Fiji to create multiple MIPs from the same Z-stack, including only optical sections that contained the dorsal nerve without neuron cell bodies in each XY position. These MIPs were adjusted to equivalent brightness and contrast and then stitched together using the Pair-wise Stitching plugin /citePreibisch2009.

### Statistical Analyses

Continuous data (Fig. 3) were analyzed using one- or two-tailed t-tests, as indicated in figure legends. Categorical data (Figs. 1–5) were analyzed in contingency tables using one- or two-tailed Fisher’s exact tests for proportionality, as indicated in figure legends. Count data (Fig. 3) were analyzed using two-tailed Mann Whitney tests.

## Supporting information

Movie S1 (related to Fig. 3)

Movie S2 (related to Fig. 3)

## AUTHOR CONTRIBUTIONS

**Patricia L. Murphy:** Conceptualization, Methodology, Validation, Formal Analysis, Investigation, Data Curation, Writing - original Draft Preparation, Writing - Review & Editing, Visualization, Funding Acquisition

**Jesse Isaacman-Beck:** Conceptualization, Methodology, Investigation, Writing –Review & Editing

**Michael Granato:** Conceptualization, Resources, Writing – Review & Editing, Supervision, Funding Acquisition

## ACKNOWLEDGEMENTS

The authors would like to thank Daniel Morales, Leah Middleton, and Lauren Walker for technical assistance. We would also like to thank Dr. Andrea Stout of the Penn Cell and Developmental Biology microscopy core and the Penn sanger sequencing core for technical support. Finally, we would like to thank PLM’s thesis committee members Dr. Wenqin Luo, Dr. Steven Scherer and Dr. Stephen DiNardo, as well as past and present members of the Granato lab, for helpful feedback on data and the manuscript.

This article was typeset in Overleaf using the Henriques Lab BioRXiv template with minor modifications.

## Supplementary Tables

**Table S1.**
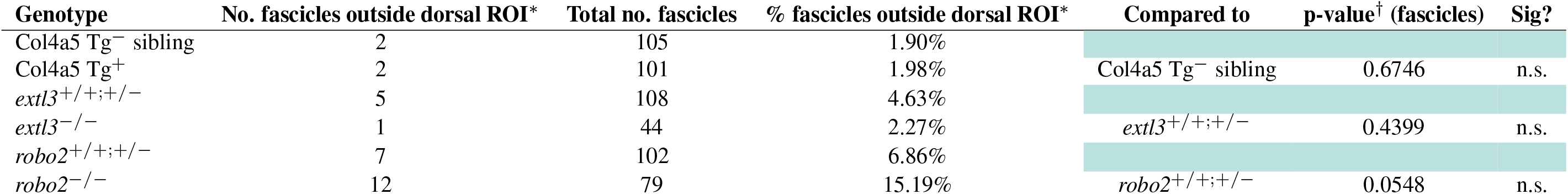
Counts of ventrolateral Tg(isl1:GFP) fascicles pre-transection (5dpf). Fascicles were counted as inside the dorsal ROI if their endpoints lay 20°-50° dorsal to the spinal cord. All statistical comparisons performed using one-tailed Fischer’s exact tests. In Sig? column, n.s. denotes “not significant.”

| Genotype | No. fascicles outside dorsal ROI* | Total no. fascicles | % fascicles outside dorsal ROI* | Compared to | p-value<sup>†</sup> (fascicles) | Sig? |
| --- | --- | --- | --- | --- | --- | --- |
| Col4a5 Tg<sup>-</sup> sibling | 2 | 105 | 1.90% |  |  |  |
| Col4a5 Tg<sup>+</sup> | 2 | 101 | 1.98% | Col4a5 Tg<sup>-</sup> sibling | 0.6746 | n.s. |
| *extl3*<sup>+/+;+/-</sup> | 5 | 108 | 4.63% |  |  |  |
| *extl3*<sup>-/-</sup> | 1 | 44 | 2.27% | *extl3*<sup>+/+;+/-</sup> | 0.4399 | n.s. |
| *robo2*<sup>+/+;+/-</sup> | 7 | 102 | 6.86% |  |  |  |
| *robo2*<sup>-/-</sup> | 12 | 79 | 15.19% | *robo2*<sup>+/+;+/-</sup> | 0.0548 | n.s. |

**Table S2.** Counts of Tg(isl1:GFP) errors at 48 hours post-transection. Fascicles were counted as inside the dorsal ROI if their endpoints lay 20°-50° dorsal to the spinal cord. All statistical comparisons were performed using one-tailed Fischer’s exact tests on contingency tables of counts of error vs. dorsal fascicles. In Sig? column, * denotes p<0.05; *** denotes p<0.001.

| Genotype | No. errors | Total no. fascicles | % errors | % errors 95CI (low) | % errors 95CI (high) | Fold change in errors | Compared to | p-value | Sig? |
| --- | --- | --- | --- | --- | --- | --- | --- | --- | --- |
| Col4a5 Tg<sup>-</sup> sibling | 29 | 96 | 30.21% | 21.93% | 40.01% |  |  |  |  |
| Col4a5 Tg<sup>+</sup> | 72 | 133 | 54.14% | 45.67% | 62.37% | +1.79 | Col4a5 Tg<sup>-</sup> sibling | 0.0002 | *** |
| *emphextl3*<sup>+/+;+/-</sup> | 31 | 98 | 31.63% | 41.38% | 23.27% |  |  |  |  |
| *extl3*<sup>-/-</sup> | 30 | 58 | 51.72% | 64.07% | 39.16% | +1.58 | *extl3*<sup>+/+;+/-</sup> | 0.0105 | * |
| *robo2*<sup>+/+;+/-</sup> | 37 | 108 | 34.26% | 25.99% | 43.61% |  |  |  |  |
| *robo2*<sup>-/-</sup> | 49 | 97 | 50.52% | 40.74% | 60.25% | +1.47 | *robo2*<sup>+/+;+/-</sup> | 0.0134 | * |

**Table S3.**
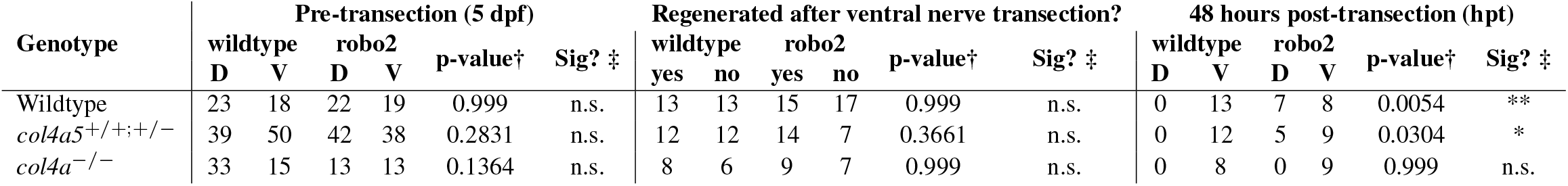
Counts of nerves with mKate^+^ labeled fascicles pre- and post-transection. Pre-transection, nerves were counted in columns labeled “wildtype” if they contained at least one visible fascicle expressing *Tg(mnx1:mKate, mnx1:mKate)*; nerves were counted in columns labeled “*robo2*” contained at least one fascicle expressing *Tg(mnx1:mKate; mnx1:robo2)*. After transection, nerves were assigned a regeneration score of “yes” if at least one mKate^+^ fascicle regenerated at least 50um along either nerve branch. At 48hpt, nerves were scored as dorsal (D) if at least one visible mKate^+^ fascicle regenerated within the dorsal ROI (20°-50° dorsal to the spinal cord). Nerves were scored as ventral (V) if there were no visible mKate^+^ fascicles within the dorsal ROI. Two-tailed Fischer’s exact tests were used to compare wildtype and robo2 conditions for each genotype pre-transection (5dpf) and to compare rates of regeneration after ventral nerve transection in wildtype and robo2 conditions for each genotype. One-tailed Fischer’s exact tests were used to compare wildtype and robo2 conditions for each genotype 48 hours post-transection. In Sig? column, * denotes p<0.05; ** denotes p<0.01, n.s. denotes “not significant.”

## Supplementary Figures

**Fig. S1.**
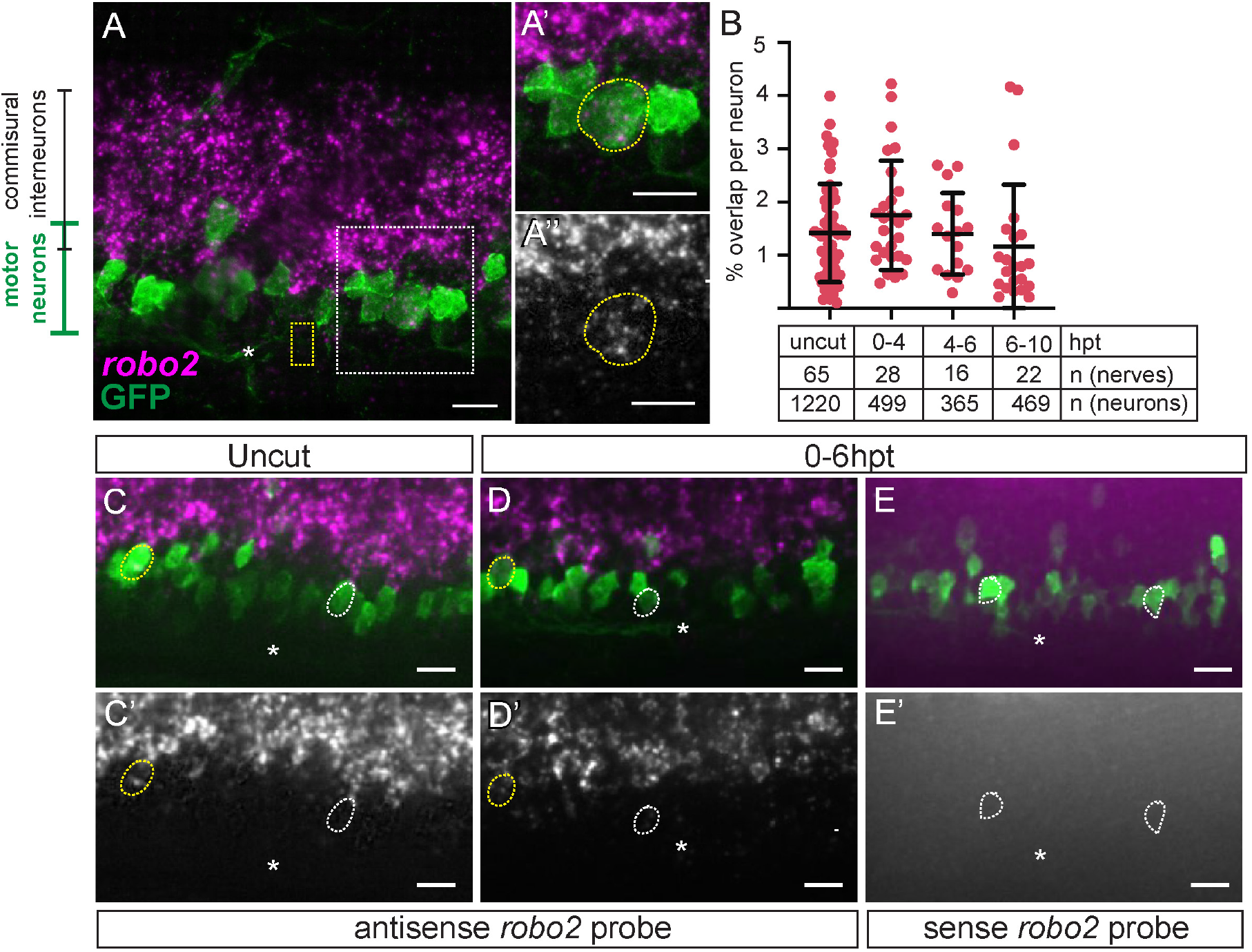
Robo2 is expressed in isl1+ motor neurons before and after transection. (**A**) Representative image of *Tg(isl1:GFP)* motor neurons in spinal cord 8-10 hpt stained with *robo2* antisense ISH probe (magenta) and GFP antibody (green). Image shown is maximum Z-projection of 24 optical sections (10um), 63X. Dashed yellow box, transection site. Dashed white box, area enlarged 1.5X in (**A’**) and (**A”**) showing a single optical section (0.41um) with one *Tg(isl1:GFP)* motor neuron (green, outlined with yellow dashed line) expressing *robo2* mRNA (magenta) merged (A’) and alone (gray) (A”). (**B**) Quantification of colocalization of *robo2* ISH probe with *Tg(isl1:GFP)* neurons at 0-10 hpt. Each dot represents the fraction of *robo2* expression in a single motor pool that colocalized with GFP, normalized to the number of labeled motor neurons. Mean % overlap per neuron was compared across timepoints using one-way ANOVA (p = 0.2138). (**C-E**) Representative images of *Tg(isl1:GFP)* motor neurons stained with *robo2* antisense ISH probe (C-D) or sense ISH probe control (E) (magenta) and GFP antibody (green) (merged). (**C’-E’**) ISH probe of corresponding image alone (gray). Images shown are single optical sections (1um), 40X. of nerves untransected (C) and at 0-6 hpt (D-E). White asterisks, motor exit points (MEPs). Yellow dashed lines outline cell bodies with expression, white dashed lines outline cell bodies without expression. Scale bars 10um.

**Fig. S2.**
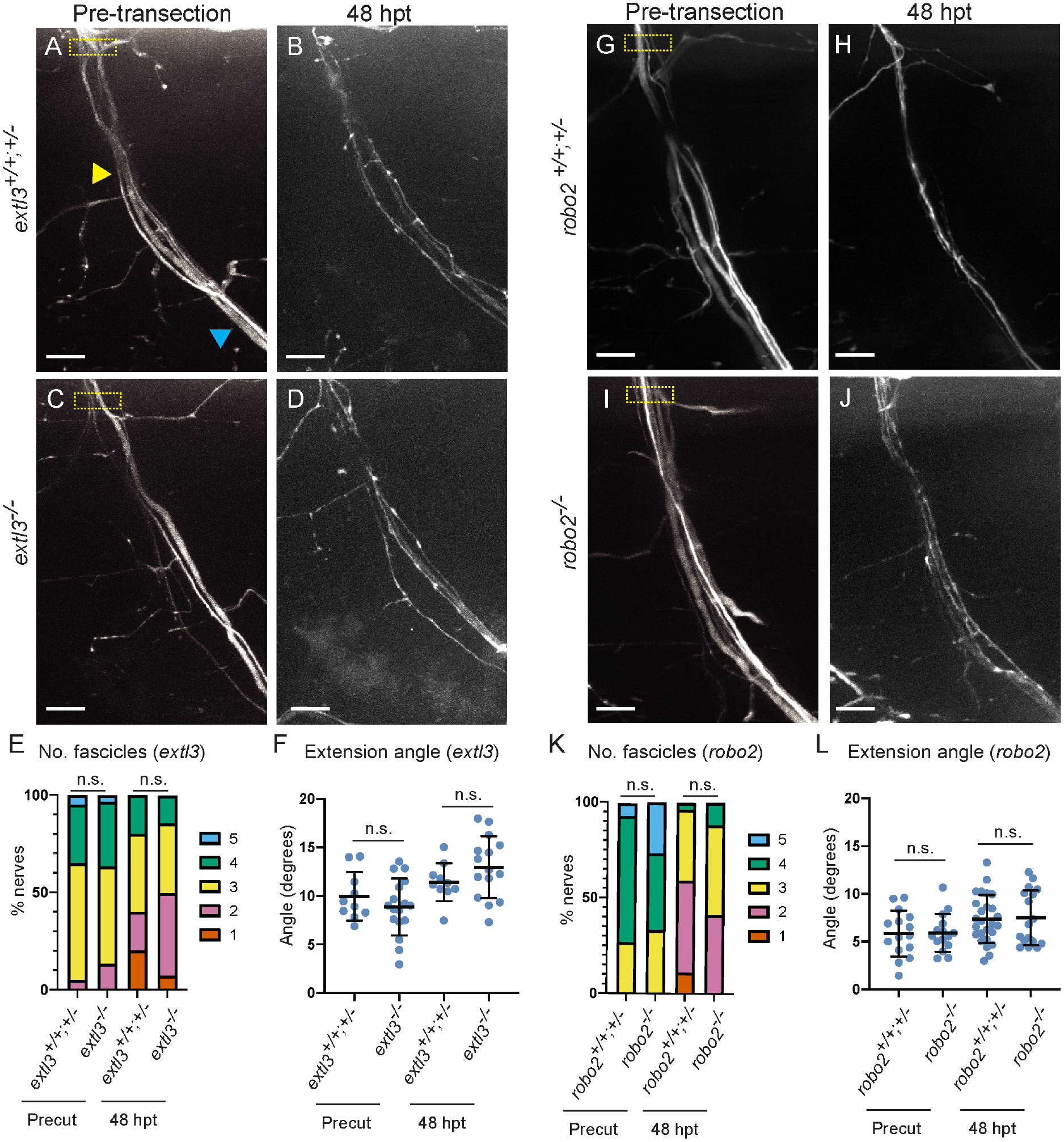
Slit-Robo signaling is dispensable for ventral nerve branch development and regeneration. Top: Representative images of *Tg(hb9:GFP)* ventral nerves at 5dpf and 48hpt, respectively, in *extl3* siblings (**A**, **B**) *extl3*^−/−^ (**C**, **D**), *robo2* wildtype siblings (**G**, **H**), *robo2*^−/−^ (**I**, **J**). Motor exit point (MEP) is just dorsal to the top of imaging frame shown. Arrowheads in (A) mark 50um (yellow arrowhead) and 100um (blue arrowhead) from the MEP. Dashed yellow boxes denote transection site (10-15um ventral to MEP). Bottom: Quantification of ventral nerve fascicles at 5dpf (Precut) and 48hpt in (**E**) extl3 wildtype siblings (n = 20 nerves, 11 larvae) and *extl3*^−/−^ larvae (n = 30 nerves, 16 larvae) and (**K**) *robo2* wildtype siblings (n = 10 nerves, 5 larvae) and *robo2*^−/−^ larvae (n = 17 nerves, 9 larvae). Nerves were scored by counting the number of discrete fascicles discernible 50um from the MEP (yellow arrowhead in A). Genotypes were compared using two-tailed t-tests (for *extl3*: Precut, p = 0.6875 ; 48hpt, p = 0.9167 for *robo2*: Precut p = 0.6009 ; 48hpt, p = 0.1003 Angle of ventral nerve extension at 5dpf and 48hpt in (F) *extl3* wildtype siblings (n = 10 nerves, 5 larvae) and *extl3*^−/−^ larvae (n = 17 nerves, 9 larvae) and (L) *robo2* siblings (n = 26 nerves, 10 larvae) and *robo2*^−/−^ larvae (n = 17 nerves, 7 larvae). Extension angle was calculated as the difference between the angle of the nerve between 50um from the MEP (yellow arrowhead in A) and 100um from MEP (blue arrowhead in A). Each dot represents one nerve. Genotypes were compared using two-tailed t-tests (for *extl3*: Precut, p = 0.3403 ; 48hpt, p = 0.1900; for *robo2*: Precut p = 0.9424 ; 48hpt, p = 0.8731).

**Fig. S3.**
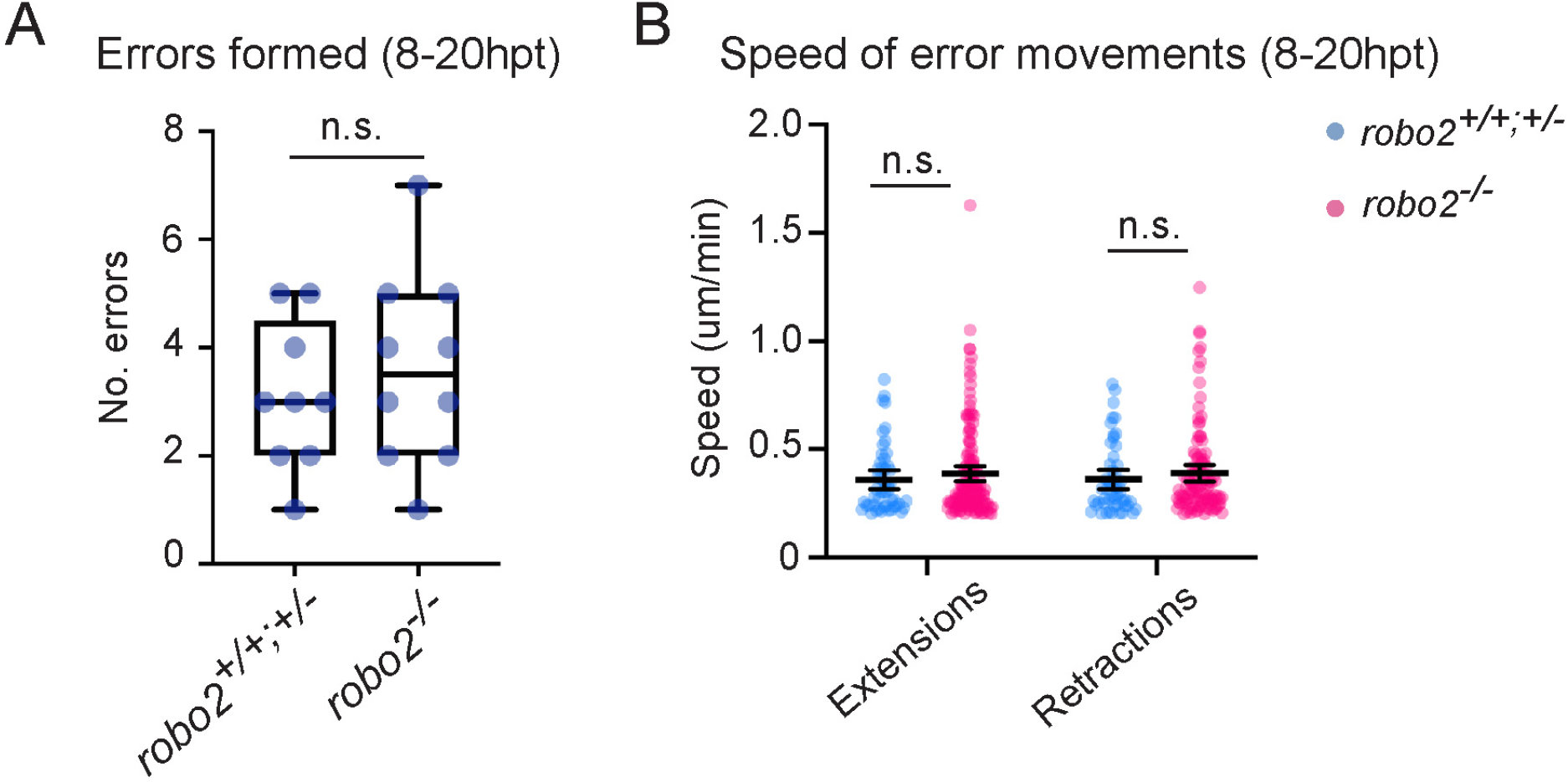
Error formation and regenerating fascicle growth dynamics are unaffected in *robo2 mutant*. (**A**) Number of errors formed 8-20hpt in wildtype siblings (*robo2*+*/*+;^+/−^, n = 9 nerves) and *robo2*^−/−^ larvae (n = 10 nerves). Each dot represents one nerve. Ranks between genotypes were compared using two-tailed Mann Whitney test (p = 0.6238) (**B**) Error extension and retraction speed calculated as the absolute value of error movement velocity in um/min. Extension and retraction events were examined in 10 min intervals and classified as extensions when there was a ≥1um increase in error length measured from the motor exit point (MEP); movements were classified as retractions when ≥1um decrease in error length measured from the MEP. Each dot represents one extension or retraction event (for siblings n = 23 errors underwent 51 extension events and 51 retraction events; for *robo2*^−/−^, n = 34 errors underwent 139 extension events and 109 retraction events). Line, mean; error bars, 95% confidence intervals. Genotypes were compared using two-tailed t-tests (for extensions, p = 0.3849; for retractions, p = 0.3853). n.s. denotes “not significant.”

**Fig. S4.**
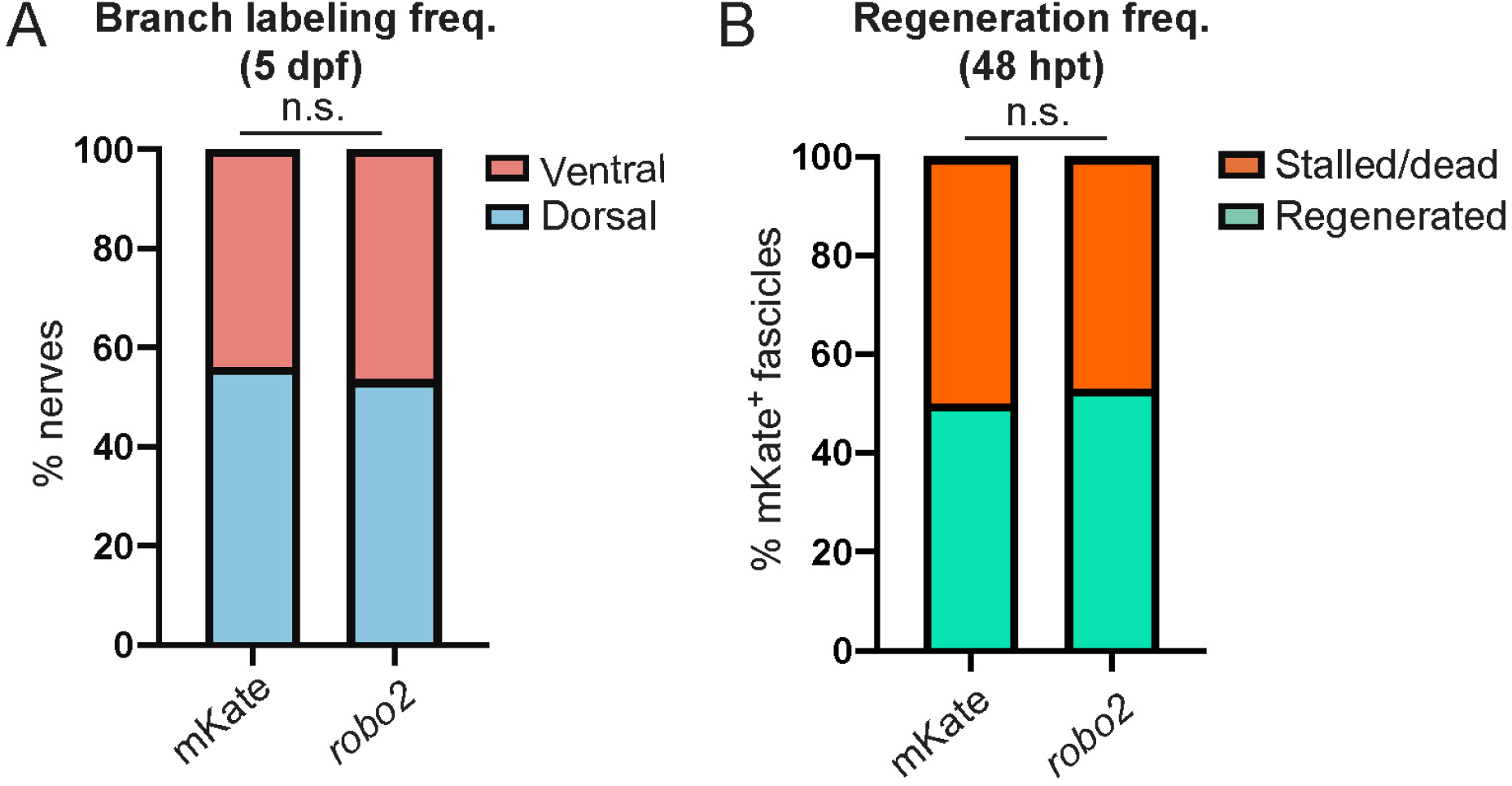
*Robo2* does not affect developmental branch-selectivity or regeneration capacity of spinal motor axons. (**A**) Graph of developmental branch-selectivity of sparsely labeled fascicles in *Tg(hb9:GFP)* larvae, showing the percent of spinal motor nerves in which any fascicles expressing *Tg(mnx1:mKate; mnx1:mKate)* (mKate in graph) or *Tg(mnx1:mKate; mnx1:robo2)* (robo2 in graph) developed along the dorsal nerve branch (dorsal) or along the ventral branch exclusively (ventral) at 5dpf (for *Tg(mnx1:mKate; mnx1:mKate)*, n = 41 nerves; for *Tg(mnx1:mKate; mnx1:robo2)*, n = 41 nerves). Two-tailed Fischer’s exact test was used to compare proportions of nerve counts in each category (p = 0.999). (**B**) Graph of regeneration frequency of fascicles showing the percent of spinal motor nerves in which fascicles expressing *Tg(mnx1:mKate; mnx1:mKate)* (mKate in graph) or *Tg(mnx1:mKate; mnx1:robo2)* (robo2 in graph) regenerated at least 50um along either nerve branch (Regenerated) or failed to regenerate (Stalled/dead) at 48 hpt (for *Tg(mnx1:mKate; mnx1:mKate)*, n = 26 nerves; for *Tg(mnx1:mKate; mnx1:robo2)*, n = 32 nerves). Two-tailed Fischer’s exact test was used to compare proportions of nerve counts in each category (p = 0.999). n.s. denotes “not significant.”

**Fig. S5.**
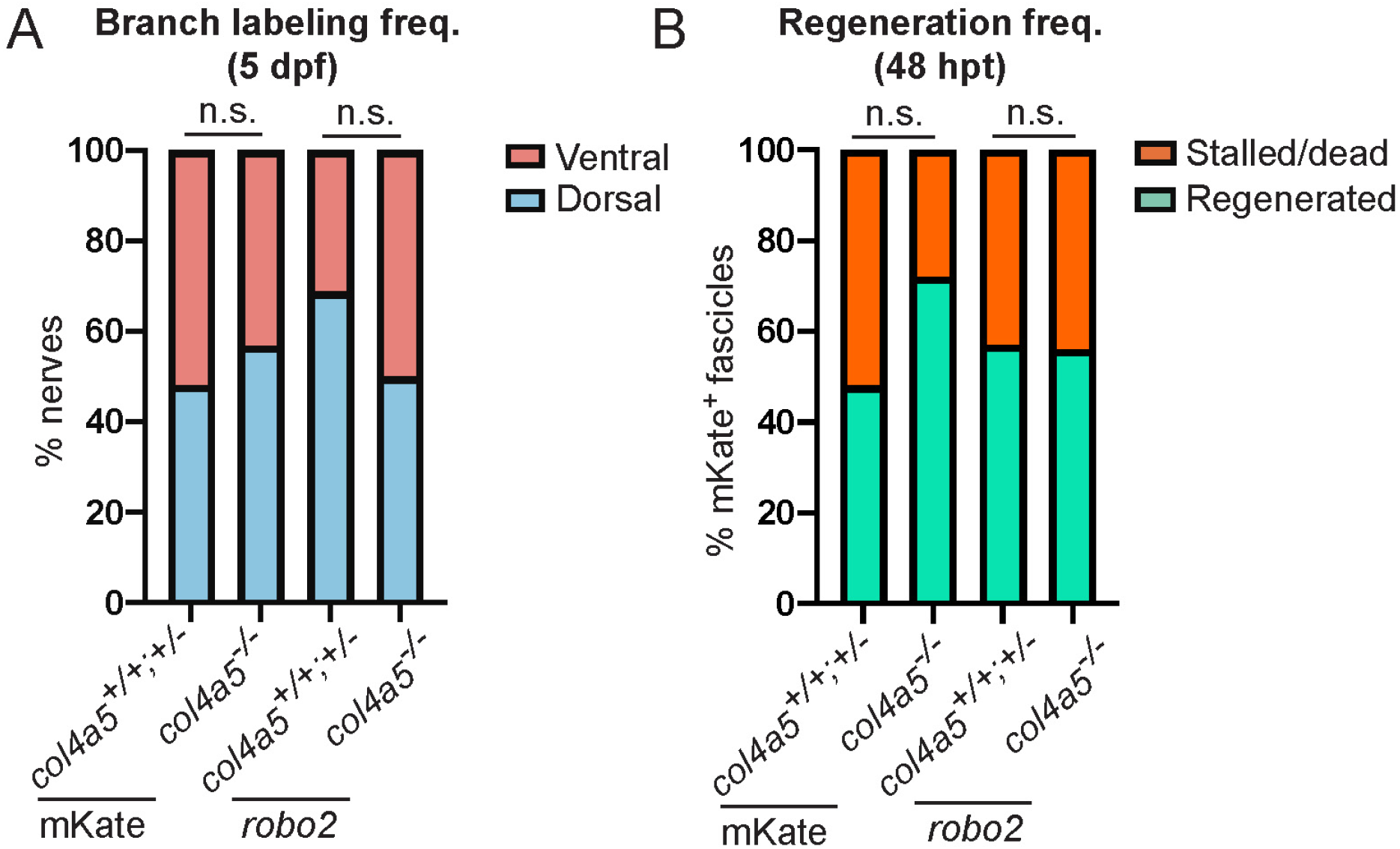
*Robo2* does not affect developmental branch-selectivity or regeneration capacity of spinal motor axons in *col4a5*^++^;^+/−^ (wildtype siblings) and *col4a5*^−/−^. (**A**) Graph of developmental branch-selectivity of sparsely labeled fascicles in in *col4a5*+*/*+;^+/−^ (wildtype siblings) and *col4a5−/− Tg(hb9:GFP)* larvae, showing the percent of spinal motor nerves in which any fascicles expressing *Tg(mnx1:mKate; mnx1:mKate)* (mKate in graph) or *Tg(mnx1:mKate; mnx1:robo2)* (robo2 in graph) developed along the dorsal nerve branch (dorsal) or along the ventral branch exclusively (ventral) at 5dpf (for wildtype siblings: *Tg(mnx1:mKate; mnx1:mKate)*, n = 89 nerves; for *Tg(mnx1:mKate; mnx1:robo2)*, n = 80 nerves; for *col4a5*^−/−^ larvae *Tg(mnx1:mKate; mnx1:mKate)*, n = 48 nerves; for *Tg(mnx1:mKate; mnx1:robo2)*, n = 26 nerves). Two-tailed Fischer’s exact test was used to compare proportions of nerve counts in each category (for siblings, p = 0.2831; for *col4a5*^−/−^ larvae, p = 0.1364). (B) Graph of regeneration frequency of fascicles showing the percent of spinal motor nerves in *col4a5*^+/+^;^+/−^ (wildtype siblings) and *col4a5−/− Tg(hb9:GFP)* larvae in which fascicles expressing *Tg(mnx1:mKate; mnx1:mKate)* (mKate in graph) or *Tg(mnx1:mKate; mnx1:robo2)* (robo2 in graph) regenerated at least 50um along either nerve branch (Regenerated) or failed to regenerate (Stalled/dead) at 48 hpt (for wildtype siblings: *Tg(mnx1:mKate; mnx1:mKate)*, n = 24 nerves; for *Tg(mnx1:mKate; mnx1:robo2)*, n = 21 nerves; for *col4a5*^−/−^ larvae *Tg(mnx1:mKate; mnx1:mKate)*, n = 14 nerves; for *Tg(mnx1:mKate; mnx1:robo2)*, n = 16 nerves). Two-tailed Fischer’s exact test was used to compare proportions of nerve counts in each category (for siblings, p = 0.3661; for *col4a5*^−/−^ larvae, p = 0.999). n.s. denotes “not significant.”

**Fig. S6.**
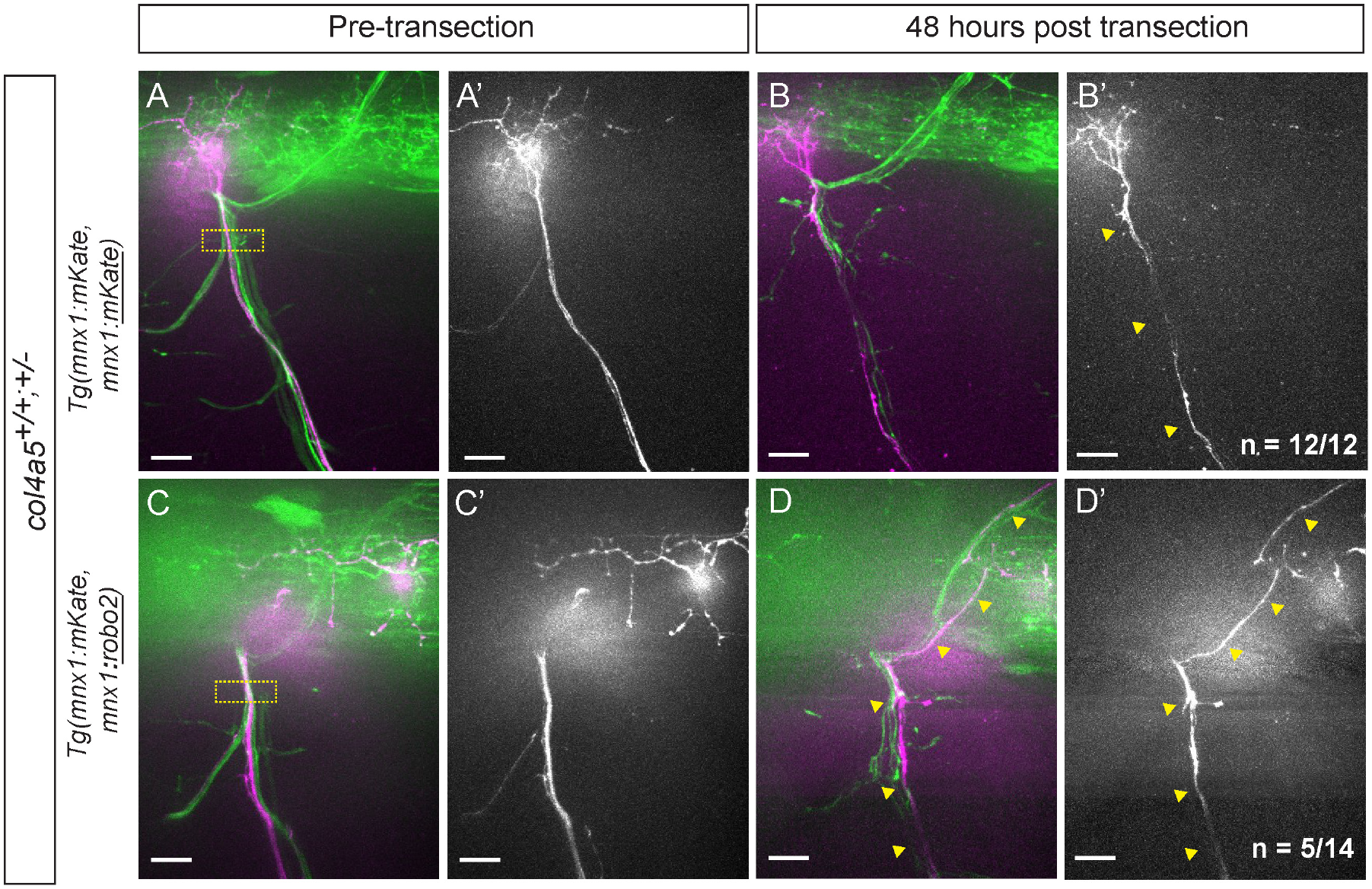
*robo2* is sufficient to promote dorsal branch-selection by regenerating axons in *col4a5*^+/+^;^+/−^ larvae (wildtype siblings). Representative images of *Tg(hb9:GFP)* (green) nerves in *col4a5*^+/−^ (wildtype sibling) larvae with fascicles expressing transient *Tg(mnx1:mKate, mnx1:mKate)* (**A-B**) or *Tg(mnx1:mKate, mnx1:robo2)* (**C-D**) (magenta) in small subsets of ventrally projecting motor axons. Merged GFP and mKate images shown at 5dpf (A, C) and 48hpt (B, D). mKate channel is shown alone at 5dpf (**A’**, **C’**) and 48hpt (**B’**, textbfD’). In larvae expressing *Tg(mnx1:mKate, mnx1:mKate)*, n = 12/12 nerves had only ventral regrowth of mKate^+^ fascicles, like in the example shown. In larvae expressing *Tg(mnx1:mKate, mnx1:robo2)*, n = 5/14 nerves had dorsal regrowth of mKate^+^ fascicles, like in the example shown. Proportions of ventral regrowth were compared between conditions using one-tailed Fischer’s exact test (p = 0.0304). Images were processed as described in Materials and Methods. Dashed yellow boxes, transection site. Scale bars, 10um.

## Legends for Supplementary Movies

**Movie S1. Dorsal nerve fascicle regeneration dynamics in wildtype sibling, related to Figure 3.** Representative movie of regenerating axons of dorsal nerve in *robo2*^+/−^ larvae imaged *in vivo* using *Tg(isl1:GFP)*. The movie begins 8 hpt, and images were taken every 10 minutes, as indicated by time counter (white, bottom right), for 12 hours. Cumulative error count (magenta, bottom right) is written as a fraction of [number errors corrected/over number errors formed]. Error fascicle movements are marked with magenta arrowheads. Movements of fascicle growing along the dorsal path are marked with green arrowheads. Images were processed as described in Materials and Methods. Scale bar is 10um.

**Movie S2. Dorsal nerve fascicle regeneration dynamics in *robo2* mutant, related to Figure 3.** Representative movie of regenerating axons of dorsal nerve in *robo2*^−/−^ larvae imaged *in vivo* using *Tg(isl1:GFP)*. The movie begins 8 hpt, and images were taken every 10 minutes, as indicated by time counter (white, bottom right), for 12 hours. Cumulative error count (magenta, bottom right) is written as a fraction of [number errors corrected/over number errors formed]. Error fascicle movements are marked with magenta arrowheads. Movements of fascicle growing along the dorsal path are marked with green arrowheads. Images were processed as described in Materials and Methods. Scale bar is 10um.

